# Repurposed endogenous virus-like vesicles mediate dendritic cell long-range antigen presentation and T cell activation for enhanced cancer vaccination

**DOI:** 10.1101/2025.11.23.690059

**Authors:** Wenchao Gu, Ruoxin Li, Hayleigh Goodrich, Dorin Artzi, Simian Cai, Aixin Shi, Irene Tsai, Guoting Qin, Ann Li, Evelyn Goldwasser, Nadine Elkasri, Sijin Luozhong, Shinji Kamada, Yizhou Wu, Vaibhav Upadhayay, Yu Zhao, Yating Yang, Justin Lau, Ella Sultan, Alexei Wirganowicz, Chenzhi Cai, Qiuming Yu, Shaoyi Jiang

## Abstract

Dendritic cells (DCs) release extracellular vesicles (DEVs) that amplify antigen presentation while incorporating patient-derived, rapidly evolving antigens. By enriching peptide-MHC and co-stimulatory ligands orders-of-magnitude above donor-cell levels, DEVs emerge as potent vesicle-vaccines, although efficacy remains limited by unclear mechanisms. We show that DCs repurpose viral components to generate endogenous virus-like vesicles (VLVs) that preferentially carry peptide-MHC and high-density co-stimulatory ligands, intensifying and extending antigen presentation. Upon antigen exposure, Arc partners with endogenous envelope proteins to assemble VLVs that directly engage T cells and trigger intrinsic adjuvanticity via viral mimicry. Arc⁻/⁻ DEVs failed to prime antigen-specific T cell responses, whereas Arc overexpression with its 5′-UTR stem-loop shifted DEVs toward VLVs that trafficked to lymphoid organs, drove rapid CD4-assisted priming and durable CD8-biased memory, suppressed melanoma, and prolonged survival. These reveal a viral-mimicry mechanism enabling long-range immune activation and support Arc^+^ VLVs as an antigen-agnostic vaccine for cancer immunotherapy.

**Highlights:**

- Arc^+^ VLVs intensify and extend DC antigen presentation *in vivo*
- Arc^+^ VLVs directly engage T cells and trigger viral-mimic adjuvanticity
- Arc^⁻/⁻^ DEVs fail to prime antigen-specific T cell responses
- Engineered Arc^+^ VLVs drive durable CD8-biased memory and tumor control
- Arc^+^ VLV vaccination provides long-range and cross-tumor protection *in vivo*

## Introduction

Dendritic cell (DC), recognized as one of the most potent antigen-presenting cells (APCs) ^1^, have emerged as a promising platform in immunotherapy. DC vaccines utilize the inherent ability of these cells to initiate and modulate immune responses. By loading DCs with disease-specific antigens and subsequently reintroducing them into the patient, they can potentially direct the immune system to recognize and combat disease cells, orchestrating a highly personalized immune response. This approach offers specificity in targeting the individual’s disease and the flexibility to present multiple disease-associated antigens, as well as the ability to accommodate mutating antigens, which is critical in cancers and chronic infectious diseases where antigenic variation is a challenge. DC vaccines can stimulate both helper and cytotoxic T cells and hold the potential for inducing long-lasting immune memory, providing ongoing protection against disease recurrence. Moreover, DC vaccines can be effectively combined with other therapies, potentially enhancing overall treatment effectiveness. However, the development of patient-specific DC vaccines faces several challenges ^2–4^, and achieving strong and sustained T cell responses remains challenging. Additionally, DC vaccines sometimes face difficulty in stimulating broad immune responses, and variability in vaccine preparation and delivery has led to inconsistent efficacy, underscoring the need for refined strategies that enhance antigen presentation and prolong immune activation.

Beyond DCs themselves, their extracellular vesicles (EVs) play a crucial role in antigen presentation, especially in facilitating long-range communication between the tumor microenvironment (TME) and lymphoid organs. DC-derived EVs (DEVs) traffic from tumor-infiltrating DCs to distant sites and engage immune cells, reflecting their parental profile by activating CD8⁺ and CD4⁺ T cells through both direct and indirect routes ^5–7^. Critically, DEVs display high densities of peptide-MHC I/II together with co-stimulatory/adhesion molecules (e.g., CD80/CD86, ICAM-1), creating potent antigen-presenting interfaces and enabling transfer of intact peptide-MHC to recipient APCs (cross-dressing) ^8–10^. As autologous vectors, DEVs deliver precise tumor antigens with lower off-target immunogenicity than synthetic agents, yet achieving consistent potency in patients remains challenging ^10–13^. Early clinical trials of dendritic cell-derived exosome (DEX) vaccines have demonstrated safety and tolerability, yet robust T cell activation remains inconsistent, particularly in tumors like melanoma and non-small cell lung cancer (NSCLC) ^14–18^. Limited T cell infiltration and the immunosuppressive TME hinder their efficacy, despite their potential for presenting tumor antigens. Ongoing efforts aim to refine DEVs to induce stronger and more sustained immune responses across diverse patient populations.

Activity-regulated cytoskeleton-associated protein (Arc) emerges as a fascinating regulator in immune modulation, particularly within the domain of DC antigen presentation ^19–21^. Originally found in migratory DCs within the skin, Arc plays a pivotal role in augmenting DCs’ response to inflammation. Furthermore, its critical roles in facilitating optimal T cell activation in cancers and autoimmune diseases highlight its significance ^20,21^. Intriguingly, Arc is a retrotransposon-derived gene encoding a protein that self-assembles into virus-like capsids, which subsequently exit host cells in the form of EVs ^22^. These endogenous virus-like vesicles (VLVs) are vehicles of long-range intercellular communication and molecular transfer ^22,23^. Drawing parallels to its retroviral Gag protein counterparts, Arc may engage envelope proteins (Envs) and other membrane-associated proteins in its assembly to interact with T cells and B cells ^24–26^. This leads to the hypothesis that Arc VLVs facilitate the transfer of antigenic information from DCs to T cells, enhancing the adaptive immune response. By increasing the proportions of Arc VLVs among total DEVs, we can improve their efficacy of antigen presentation, utilizing the unique properties of Arc in enhancing DC function and interaction with T cells.

This research represents an advancement in cancer vaccination by enabling antigen incorporation into engineered endogenous VLVs without requiring the identification of specific neoantigen sequences. This approach allows highly personalized cancer therapies using antigens derived from a patient’s own tumor, offering a tailored and comprehensive immune response. The system’s adaptability is crucial given the dynamic nature of cancer, where rapid mutations lead to the constant emergence of neoantigens. This method seamlessly integrates newly arising tumor-specific antigens into the vaccine formulation, using a variety of tumor-derived proteins and EVs. For instance, following tumor resection, cultured tumor organoids can secrete antigenic proteins and EVs, creating DEV vaccines for recurrence prevention. In patients with active disease, biopsies or circulating tumor-associated EVs provide additional sources of relevant antigens, allowing the vaccine to stay effective against the tumor’s evolving landscape.

Moreover, our findings unveil a previously unrecognized dimension of immunology, where DCs repurpose ancient viral components to enhance communication with T cells. This mechanism, by which DCs harness endogenous virus-like elements to achieve long-range immune activation, reveals an evolutionary strategy where viral components are integrated into host immune functions, enhancing immune surveillance and response. Proteomic analysis shows that Arc VLVs not only utilize viral machinery to engage T cells effectively but also function as natural adjuvants that strengthen immune responses while also presenting antigens more efficiently. This offers valuable insights into the potential of utilizing endogenous virus-like elements for highly biocompatible yet effective vaccine strategies. Such approaches could transform immunotherapies for a broad spectrum of conditions, including cancers, infectious, autoimmune, and neurodegenerative diseases.

## Results

To investigate the role of Arc VLVs in mediating long-range intercellular communication between immune cells, we first studied how Arc naturally responds to external stimulation in APCs, particularly in DCs and macrophages (MΦs). Extracted and differentiated from mice, primary bone marrow-derived DCs and MΦs (BM-DC/MΦs) ^23^ were exposed to various stimuli including lipopolysaccharide (LPS), ovalbumin (Ova), and lipid-based transfection reagents (liposomes/lipofectamine), selected to model innate pathogen-associated cues (LPS), antigenic processing (Ova), and the synthetic perturbations introduced by the very reagents we use for donor-cell engineering (lipid transfection), allowing assessment of Arc induction across all relevant activation contexts. Reverse transcription quantitative PCR (RT-qPCR) analysis revealed a rapid increase in *Arc* mRNA expression half an hour post-stimulation (Fig. 1A, purple, right y-axis). EVs collected 6 h post-stimulation showed increased intravesicular Arc signal by on-bead permeabilized immunofluorescence: vesicles were captured with CD81 beads, permeabilized, stained with a fluorescent anti-Arc antibody, and read on an epifluorescence plate reader (Fig. 1A, black, left y-axis). Together, these results suggest that DCs and MΦs initiate Arc expression and secrete Arc^+^ EVs in response to various stimuli, indicating a critical role of Arc following APC activation.

**Figure 1.**
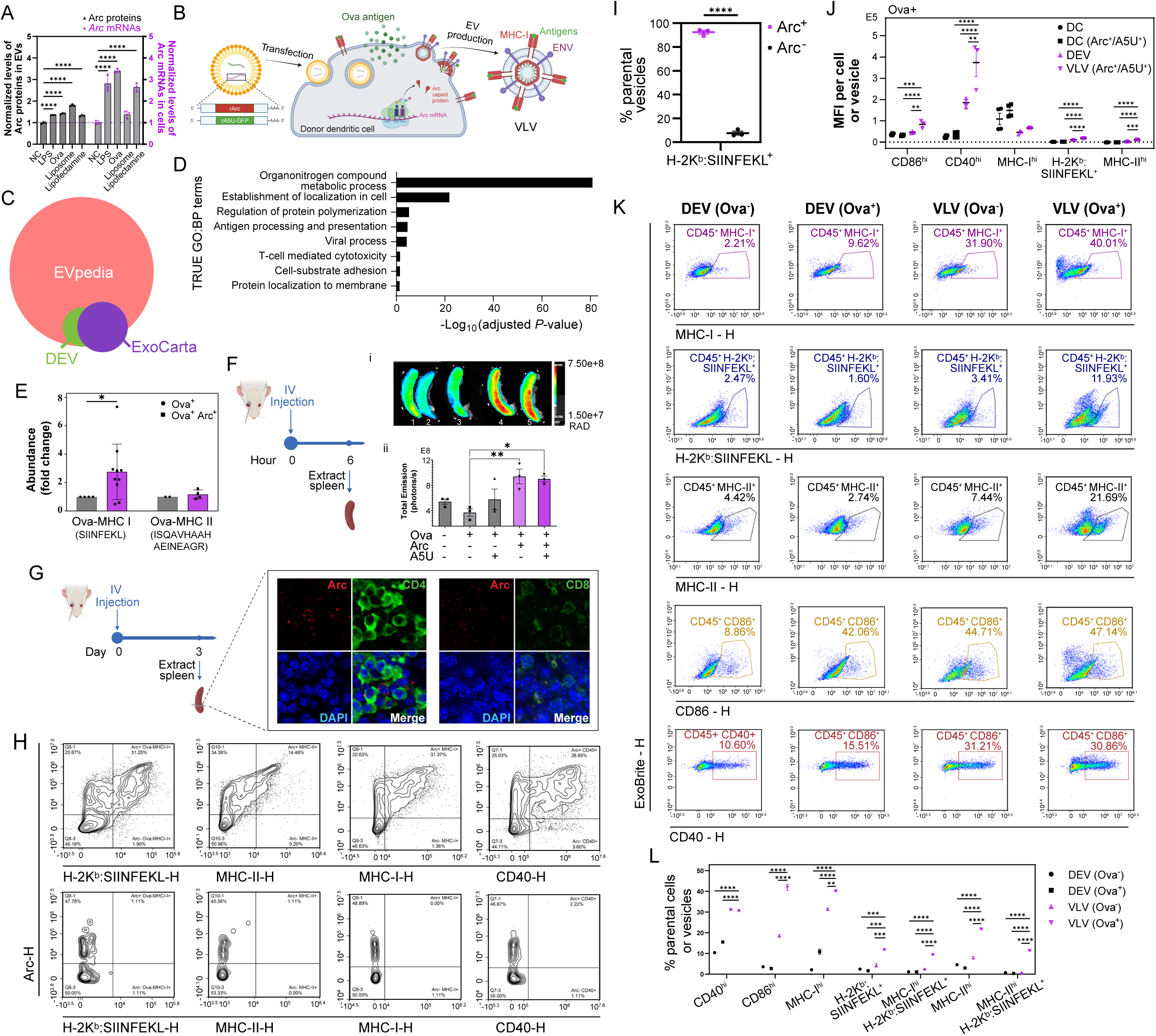
Arc VLVs respond to stimuli in APCs, present antigens, target secondary lymphoid organs, and directly interact with T cells. **(A)** Donor BM-DCs/MΦs were stimulated for 6 h with LPS, Ova, Xfect liposomes or lipofectamine. Cellular *Arc* mRNA (RT-qPCR; purple, right y-axis; normalized to NC) and EV-associated Arc protein (permeabilized anti-Arc fluorescence; black, left y-axis) are shown. Bars = mean ± SEM (*n* = 3 biological replicates). Statistics: one-way ANOVA with Dunnett’s multiple-comparisons test *vs.* NC, performed independently for each readout; **** *P* < 0.0001. **(B)** Schematic of VLV engineering: DC2.4 cells were engineered to co-express Arc and the A5U stabilizer RNA element and were subsequently pulsed with the model antigen Ova to generate Ova⁺ Arc⁺/A5U⁺ VLVs. **(C)** Venn diagram showing the overlap of proteins identified by MS with those listed in established EV protein databases (ExoCarta and EVpedia). **(D)** TRUE enriched GO:BP terms for DEVs, identified using G:Profiler. **(E)** Mass spectrometry quantification of Ova peptide presentation: A significant increase in Ova:MHC-I (SIINFEKL) abundance was observed in Arc⁺ VLVs compared with WT DEVs, whereas Ova:MHC-II (ISQAVHAAHAEINEAGR) showed no significant difference. Data are mean ± SEM from four to ten independent readings. Statistics: unpaired two-tailed t-test with Welch’s correction (P = 0.0192). **(F)** (i) IVIS imaging 6 h post-IV injection of control or Arc⁺ vesicles. (ii) Quantification showing significantly greater spleen enrichment of Arc⁺ DEVs and Arc⁺ A5U⁺ VLVs. Mean ± SEM (n = 3); *P = 0.0111, **P = 0.0071; one-way ANOVA. **(G)** Cryo-sectioned spleen images showing robust accumulation of Arc⁺ VLVs and direct binding to splenocytes (Arc shown in red; CD4/CD8 in green; DAPI in blue). **(H)** Single-vesicle EV flow cytometry (representative contour-density plots): (Top panels) Arc fluorescence (y-axis) plotted against H-2Kᵇ:SIINFEKL, MHC-II, MHC-I, or CD40 (x-axis), showing that the predominant H-2Kᵇ:SIINFEKL⁺, MHC-II⁺, MHC-I⁺, and CD40^hi^ vesicle subsets are uniquely Arc⁺. (Bottom panels) FMO controls stained for Arc only, used to define the positive gates for each target. **(I)** Quantification of Arc⁺ H-2Kᵇ:SIINFEKL⁺ vs. Arc⁻ H-2Kᵇ:SIINFEKL⁺ vesicles: Arc⁺ vesicles nearly exclusively display H-2Kᵇ:SIINFEKL. Mean ± SEM (n = 3); unpaired two-tailed t-test; ****P < 0.0001. **(J)** MFI per particle for CD86, CD40, MHC-I, H-2Kᵇ:SIINFEKL, and MHC-II measured on live singlet CD45⁺ CD11c^+^ DCs or singlet CD9⁺ CD45⁺ CD11c^+^ vesicles: DCs (*n* = 4), DEVs (*n* = 3), Arc^+^/A5U^+^ DCs (*n* = 4) and VLVs (*n* = 3). Data shown are representative of 3 independent experiments, each performed with 2-4 replicates (one replicate is a separate T75 flask of donor cells). Controls include dye-only, no-stain EV-only, detergent^+^ EV and FMOs. *P* values were determined by one-way ANOVA Dunnett’s multiple comparisons test, performed independently for each target: CD86^hi^ (\*\*\**P* = 0.0001, \*\*\*\**P* < 0.0001, \*\**P* = 0.0011); CD40^hi^ (\*\*\*\**P* < 0.0001, \*\**P* = 0.0032); H-2Kᵇ:SIINFEKL^+^ (\*\*\*\**P* < 0.0001); MHC-II^hi^ (\*\*\*\**P* < 0.0001, \*\*\**P* = 0.0002). **(K)** Representative dot plots comparing surface expression of MHC-I, H-2Kᵇ:SIINFEKL, MHC-II, CD86, and CD40 between singlet CD9⁺ CD45⁺ vesicles and parental donor DCs. **(L)** Quantification of % of parental vesicles for CD86^hi^, CD40 ^hi^, MHC-I ^hi^, H-2Kᵇ:SIINFEKL^+^, and MHC-II^hi^ across DEV (Ova⁻/Ova⁺) and VLV (Ova⁻/Ova⁺) groups. Mean ± SEM (*n* = 3). One-way ANOVA with Dunnett’s multiple-comparisons test for each marker: ****P < 0.0001; **P = 0.0018 (MHC-I^hi^); ***P = 0.0002; ***P = 0.0002; ***P = 0.0005 (H-2Kᵇ:SIINFEKL⁺).

To study the composition and functionality of DC-derived VLVs, we first genetically engineered donor DCs to produce Arc^+^ VLVs (Fig. 1B), as previously described ^23^. Overexpressing the Arc capsid protein was insufficient to significantly increase production of Arc^+^ VLVs, and we complemented the overexpression with a stabilizing RNA motif, the 5’ UTR of the Arc gene (A5U) ^23^. We used rat Arc for easy differentiation from mouse Arc in *in vivo* models and human Arc in *in vitro* models. Additionally, we incorporated a GFP reporter linked to the A5U to monitor transfection efficiency. Using liposome-mediated transfection of RNAs encoding Arc and A5U-GFP (Fig. S1A), we substantially increased engineered VLV output, routinely achieving >70% Arc⁺ vesicles among total EVs (Fig. S1B) ^23^. Single-vesicle EV flow cytometry was performed on a NovoCyte Quanteon operated at the lowest flow rate (5 μL/min), which allows reliable detection of fluorescent particles down to 100 nm. Because antibody removal is a major challenge for EV flow, we cleaned stained samples using the EXODUS H-600 automatic exosome isolation system (Exodus Bio), which employs dual-membrane nanofiltration with periodic negative pressure oscillation and ultrasonic harmonic oscillations to efficiently remove free proteins and antibodies while retaining intact vesicles. Controls included EV-only (unstained), stain-only (no EVs), fluorescence-minus-one (FMO) controls, and detergent-lysed EVs with staining to distinguish true EV signals from background.

Having established a robust system to generate and validate Arc⁺ VLVs, we next assessed their antigen-handling capacity. We added the Ova model antigen directly to the culture medium, allowing both control and engineered DCs to process and present the antigen. Control DEVs and engineered VLVs were subsequently purified using tangential flow filtration (TFF), ultrafiltration, or EXODUS-based filtration, depending on the required level of purity. Nanoparticle tracking analysis (NTA) revealed a distinct subpopulation of larger vesicles in the engineered preparations, consistent with retrovirus-like capsids budding from the plasma membrane (Fig. S1C) ^23^. Western blot analysis confirmed the expected EV marker composition of these vesicles (Fig. S1D) ^23^. Liquid chromatography-tandem mass spectrometry (LC-MS/MS) analysis provided a detailed protein profile of the vesicles, showing significant overlap with documented EV proteins from databases such as ExoCarta and EVpedia (Fig. 1C). The G:Profiler database identified several TRUE Gene Ontology Biological Process (GO:BP) terms associated with these proteins, which emphasize the vesicles’ capabilities in immune-related functions (Fig. 1D). These include organonitrogen compound metabolic process, establishment of localization in cell, regulation of protein polymerization, antigen processing and presentation, inositol biosynthetic process and protein homooligomerization, all of which are crucial for the effective transport, processing, and presentation of antigens. Other identified functions include involvement in viral processes and T cell mediated cytotoxicity, highlighting the vesicles’ potential in activating immune responses. Additionally, proteins linked to cell-substrate adhesion indicate impacts on vesicle migration and target cell interaction. Label-free MS quantification highlighted that Ova⁺ VLVs displayed coordinated enrichment of surface adhesion and budding proteins, Icam1, Alcam, CD9, and Annexin A5, relative to control DEVs, while core cytosolic cargo (Ywhae, Annexin A2, ENO1) remained unchanged (Fig. S2A). In contrast, CD63 and Lamp2 were reduced, indicating a shift away from late endosomal/exosomal content toward a more plasma-membrane/microvesicle-like profile. Functionally, this surface remodeling predicts stronger binding and uptake by target cells, consistent with enhanced presentation capacity observed in downstream assays. Furthermore, proteins involved in both MHC-I and MHC-II antigen processing pathways (Fig. S2B, i-ii), as well as membrane proteins crucial for T cell interaction and activation were upregulated in Arc^+^ vesicles (Fig. S2B, iii). Importantly, we observed substantial increase of the MHC-I Ova peptide SIINFEKL on VLVs (Fig. 1E), whereas the MHC-II Ova peptide ISQAVHAAHAEINEAGR remained comparable. These compositional changes collectively indicate that Arc⁺ vesicles acquire a spectrum of immune-modulating membrane proteins from DCs, tailored for effective long-range intercellular communication and immune activation, while preferentially enhancing MHC-I-mediated antigen presentation.

To further investigate the roles of VLVs *in vivo*, we systemically administered both control and engineered vesicles presenting Ova peptides to study their distribution and interactions with target organs and cells. Labeled with a lipid membrane dye (CellMask Deep Red, or CMDR) for tracking, equal quantities from each sample group, including EV^-^ control (PBS), Ova^-^ DEVs, Ova^+^ DEVs, Ova^+^/A5U^+^ DEVs, Ova^+^/Arc^+^ DEVs, and Ova^+^ Arc^+^/A5U^+^ VLVs (abbreviated ‘VLV’ below), were injected intravenously into C57BL/6 mice. Six hours post-injection, the spleens were harvested and analyzed using an *in vivo* imaging system (IVIS), demonstrating a substantial enrichment of Arc^+^ vesicles (Fig. 1F). This highlights the efficient trafficking of Arc^+^ VLVs to a key secondary lymphoid organ, implying their potential in effectively reaching sites critical for initiating T cell responses. Further analysis of immunohistochemically-stained spleen cryosections revealed VLVs’ binding to spleen cells (Fig. 1G & S3). Thus, we demonstrated that APCs initiate Arc expression and secrete Arc^+^ VLVs in response to various stimuli, with these vesicles showing enhanced MHC-I antigen presentation capacity and efficiently trafficking to secondary lymphoid organs to directly interact with T cells, highlighting their crucial roles in APC functionality and therapeutic relevance in immunotherapies.

Having established these *in vivo* trafficking and antigen-presentation properties, we next investigated, at single-vesicle resolution, whether antigen-primed Arc⁺ DCs and their secreted vesicles display coordinated upregulation of surface co-stimulatory and antigen-presentation molecules, reflecting enhanced donor-cell activation and vesicle immunogenicity. We performed flow cytometry profiling of both donor DCs and purified vesicles, using a parallel gating hierarchy for both EV and cell flow cytometry: time/scatter stability → singlets → (cells) live CD45⁺ CD11c⁺ DCs; (EVs) ExoBrite⁺ CD45⁺ CD11c⁺ vesicles with dye-only and unstained controls (Fig. S4). Within these gates, we quantified Arc, MHC-I, MHC-II, CD80, CD86, CD40, and H-2Kᵇ:SIINFEKL (MHC-I:OVA₂₅₇-₂₆₄). Single-vesicle EV flow cytometry revealed an interesting feature of engineered Arc⁺ VLVs: Arc expression tightly marks the immunogenic subpopulation. In representative contour-density plots, among the FMO-defined positive populations, vesicles expressing MHC-I, MHC-II, or H-2Kᵇ:SIINFEKL, as well as those with high CD40, are almost exclusively Arc⁺, with very few events falling into the Arc⁻ gate (Fig. 1H). Quantification across three independent experiments (Fig. 1I) further confirmed that Arc⁺ vesicles account for the dominant H-2Kᵇ:SIINFEKL⁺ population.

With the immunogenic Arc⁺ subset defined, we next asked how the antigen-presentation and co-stimulatory capacity of individual vesicles compares to that of their donor DCs. Even without surface area normalization, the MFI of CD86, CD40, H-2Kᵇ:SIINFEKL, and MHC-II per vesicle exceeded that of each donor cell (Fig. 1J). Given their geometry, a 150-nm EV has ∼0.071 µm² surface area compared to ∼707 µm² for a 15-µm DC (a ∼10,000-fold difference), making surface-normalized comparisons essential. We therefore defined an Enrichment Index (EI) as the relative ligand density on each vesicle versus a donor DC, calculated as described in the Table 1 legend. Applying this surface-normalized metric, DC-derived vesicles are massively enriched for antigen-presentation and co-stimulatory displays compared to the donor cell membrane (Table 1). To capture effective output, we additionally calculated a Practical Delivery Index (PDI), incorporating secretion rate (∼1,000 EVs per DC per day, based on NTA of every DEV/VLV batch). PDI revealed that VLV-mediated delivery of MHC-I/II and co-stimulatory ligands is amplified by ∼10²-10⁶-fold relative to a single DC’s static surface display (Table 2).

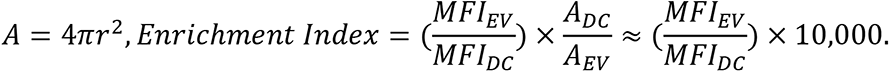

**Table 1.** EI of co-stimulatory and antigen-presentation molecules on EVs relative to donor DCs. EI was calculated from the ratio of MFI values normalized to surface area, assuming spherical geometry for a 150-nm vesicle (*r_EV_* = 0.075μ*m*) and a 15-µm DC (*r_DC_* = 7.5μ*m*):

| Marker | Control DEVs vs. mock DCs | Engineered VLVs vs. transfected DCs |
| --- | --- | --- |
| **CD86** | $1.25 \times 10^4$ | $2.63 \times 10^4$ |
| **CD40** | $7.21 \times 10^4$ | $1.09 \times 10^5$ |
| **MHC-I** | $4.21 \times 10^3$ | $4.39 \times 10^3$ |
| **SIINFEKL:H-2K<sup>b</sup>** | $9.35 \times 10^4$ | $9.61 \times 10^4$ |
| **MHC-II** | $4.44 \times 10^7$ | $9.85 \times 10^7$ |

**Table 2.**
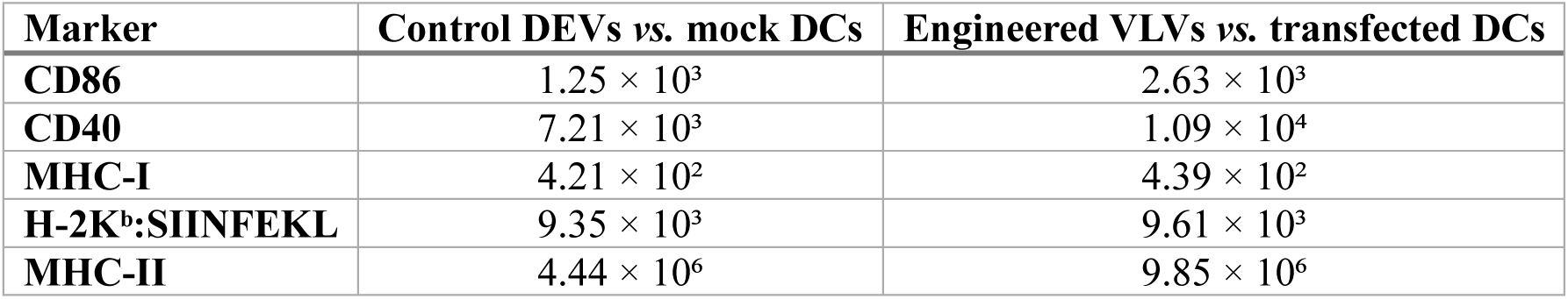
PDI of co-stimulatory and antigen-presentation molecules on EVs relative to donor DCs. PDI reflects the total delivery power of vesicles compared to their donor cells, how many more antigen-presenting or co-stimulatory molecules EVs collectively deliver to T cells.

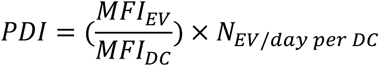

with *N*_EV/day per DC_ ≈ 10^3^.

To complement these per-surface and per-vesicle density measurements, we also examined the composition of the vesicle population. Specifically, we quantified the fractions of vesicles that were co-stimulatory^high^ (CD40^hi^/CD86^hi^), MHC-I^hi^, MHC-II^hi^, and H-2K^b^:SIINFEKL⁺. Quantitatively, VLVs contained higher fractions of CD40^hi^ (1.97×), CD86^hi^ (15.4×), MHC-I^hi^ (3.7×), MHC-II^hi^ (7.3×), and H-2K^b^:SIINFEKL⁺ (7.1×) vesicles than DEVs (Fig. 1K-L), mirroring the proteomic enrichment of adhesion and budding proteins (Fig. 1E). Together, flow cytometry analyses show that DC vesiculation dramatically expands antigen-presentation and co-stimulatory capacity beyond what can be achieved at the cell surface alone. Arc marks the dominant immunogenic vesicle subset, which preferentially carries H-2Kᵇ:SIINFEKL, MHC-I/II, and co-stimulatory ligands at high density. Arc⁺ VLVs have both higher per-vesicle ligand density and higher population-level frequencies of peptide-MHC^+^ and co-stimulatory^hi^ vesicles than conventional DEVs. Thus, Arc⁺ VLVs emerge as the dominant immunogenic vesicle type and potent amplifiers of DC-mediated immunity.

Systemic administration of Arc⁺ VLVs triggered rapid transcriptional and cellular immune activation. RT-qPCR at 24 h post-injection showed broad immune-gene upregulation in lymph nodes (pooled six cervical and four axillary/brachial) relative to the Ova⁻ DEV group, for IFN-γ, IL-2, IL-4, IL-10, and CD86 (Fig. 2A). This indicates activated APC co-stimulation (CD86) and a balanced, CTL-permissive cytokine milieu (IFN-γ/IL-2 for rapid CD8⁺ priming, IL-4 for humoral support, and IL-10 preventing overactivation) in contrast to the spleen, where no obvious changes were detected at this time point (Fig. S5). We profiled an early 24-h time point because the mechanism of action is immediate: our VLVs display peptide-MHC and co-stimulatory ligands on their surface, enabling direct engagement of T cells *in vivo*. Antigen uptake and processing occur *ex vivo* in the donor DCs prior to vesicle isolation, so no additional *in vivo* APC uptake/processing step is required. Accordingly, an early window is the most appropriate to capture the initial lymph-node transcriptional program and rapid T cell activation driven by VLVs. Consistent with this mechanism, at day 1 post-i.v. administration of Arc⁺ VLVs from Ova-primed DC2.4 donors, we observed a significant increase in CD3⁺ and CD4⁺ T cell proliferation versus PBS, accompanied by a transient reduction in CD8⁺ T cell frequency (Fig. 2B i-iii). By day 3, both total CD8⁺ T cells and CD8⁺CD44^hi^CD62L^hi^ subsets were significantly increased (Fig. 2Biii, S6), while CD4⁺ and CD4⁺CD44^hi^CD62L^hi^ frequencies declined (Fig. S6). This kinetic shift is consistent with rapid CD4-assisted priming that matures within 72 h into a CD8-biased, MHC-I-skewed response. In summary, DC-derived VLVs present cognate peptide-MHC with co-stimulatory ligands directly to T cells *in vivo*, inducing a balanced transcriptional program and a rapid CD4-assisted priming that transitions to CD8-skewed expansion.

**Figure 2.**
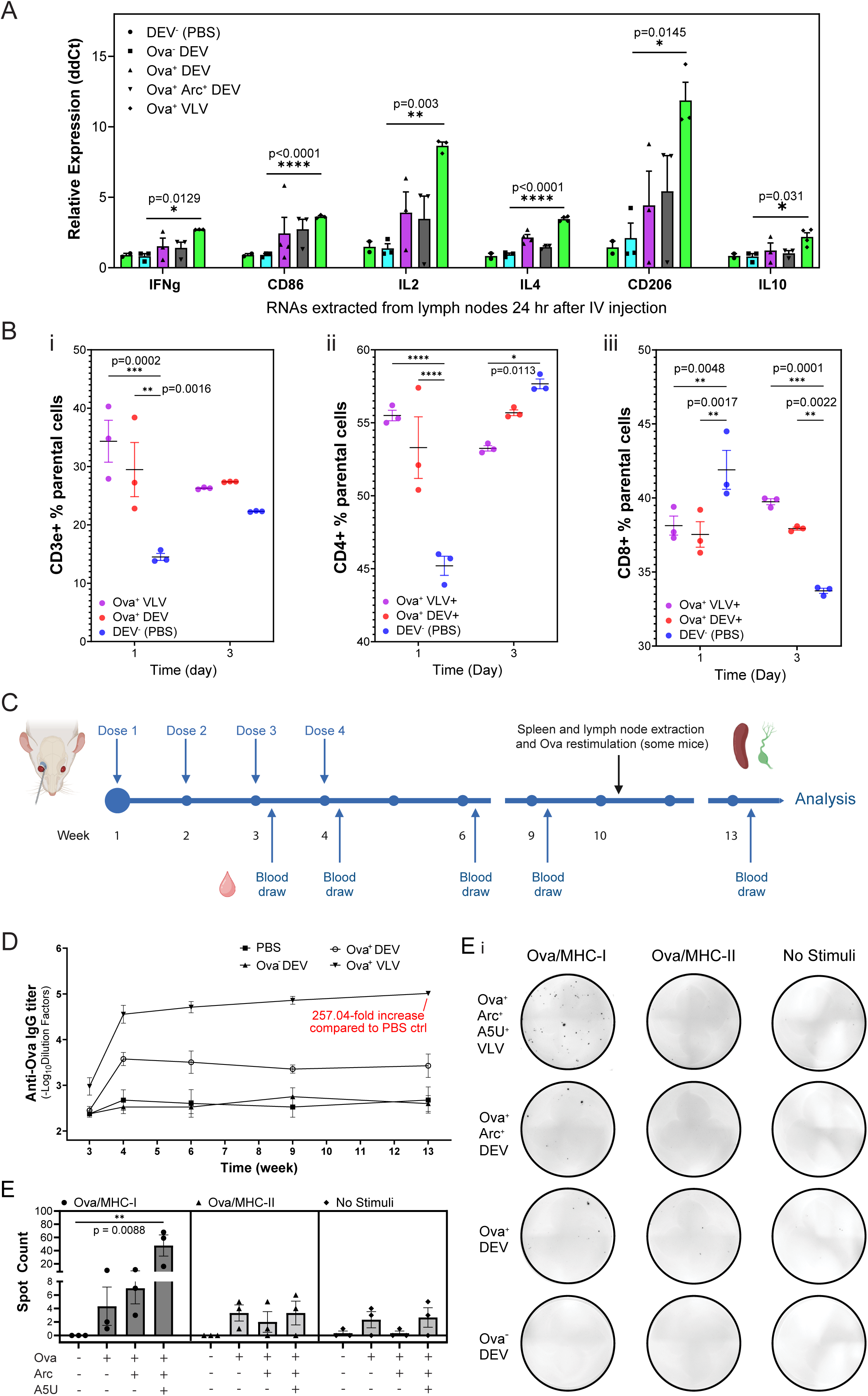
Presenting a model antigen Ova, Arc^+^ VLVs induce robust, balanced cellular and sustained humoral immune responses. In panels A-C, mice were injected with antigen^+^ control and engineered vesicles one to three days before the collection of organs in various independent experiments (*n* ≥ 3). **(A)** RT-qPCR of immune-related gene expression in lymph nodes (2 superficial cervical, 2 mandibular, 2 axillary, 2 brachial, and 2 inguinal per mouse) 24 h after IV injection. Upregulation of IFN-γ, CD86, IL-2, IL-4, CD206, and IL-10 was observed in Arc⁺ groups (Ova⁺ Arc⁺ DEVs and Ova⁺ VLVs) compared with Ova⁻ and Ova⁺ DEV controls. Mean ± SEM (n = 2-4). One-way ANOVA with Dunnett’s multiple-comparisons test vs. control; P values are shown in the panel. **(B)** Flow cytometry of splenocytes at days 1 and 3 after injection. (i) CD3⁺, (ii) CD4⁺, and (iii) CD8⁺ T cells shown as the percentage of parental cells. On day 1, mice receiving Ova⁺ VLVs showed a significant increase in the percentages of CD3⁺ and CD4⁺ T cells and a significant decrease in the percentage of CD8⁺ T cells. By day 3, the percentage of CD4⁺ T cells had decreased, whereas the percentage of CD8⁺ T cells had increased in the VLV group. One-way ANOVA with Dunnett’s multiple-comparisons test vs. PBS control; P values are shown in the panels. **(C)** Dosing and sampling schedule for the long-term study: mice received four weekly doses of control and experimental vesicles, with blood collected at the indicated time points for antibody-titer ELISA, and spleen and lymph nodes harvested after week 13 for ELISpot restimulation assays using Ova:MHC-I and Ova:MHC-II peptides. **(D)** Anti-Ova IgG titers measured by ELISA in serum from mice vaccinated with PBS, Ova⁻ DEVs, Ova⁺ DEVs, or Ova⁺ VLVs. Mean ± SEM (*n* = 4). Ova⁺ VLVs elicited significantly higher IgG titers than PBS at week 4 (**P = 0.0040), week 6 (*P = 0.0153), week 9 (**P = 0.0037), and week 13 (*P = 0.0192) by one-way ANOVA with Dunnett’s multiple-comparisons test vs. PBS. At week 13, the Ova⁺ Arc⁺ DEV group showed a 257.04-fold increase in titer relative to PBS. **(E)** ELISpot recall response after booster: some mice received a booster dose 2 months after the initial immunization, and ELISpot was performed 3 days later. (i) Representative ELISpot wells from splenocytes of mice vaccinated with Ova⁺ VLVs, Ova⁺ Arc⁺ DEVs, or Ova⁺ DEVs, restimulated *ex vivo* with Ova:MHC-I peptide, Ova:MHC-II peptide, or no stimulus. (ii) Quantification of spot counts under the indicated restimulation conditions for the different vaccine formulations (Ova / Arc / A5U combinations). Mean ± SEM (*n* = 3). Statistics: one-way ANOVA with P values indicated in the panel.

We next explored whether multiple administrations of Ova^+^ VLVs could generate a sustained humoral immune response. When administered weekly over four weeks (Fig. 2C), VLV immunization elicited a strong and sustained IgG response, indicative of robust activation of adaptive immunity. Antibody titers were significantly higher than those induced by control groups, with a 257-fold increase compared to the PBS control at week 13, demonstrating superior immunogenic potency of VLVs (Fig. 2D). Additionally, Ova⁺ VLVs triggered durable, MHC-I/Ova (SIINFEKL)-specific IFN-γ ELISpot responses with a clear booster recall two months after the initial four weekly doses, indicating long-term, MHC-I-biased memory (Fig. 2E i-ii). The H-2Kᵇ:SIINFEKL specificity reflects effective class-I presentation to CD8⁺ T cells, and the IFN-γ signal is consistent with effector/memory CD8⁺ activation, crucial in cancer vaccine strategies. In summary, repeated VLV dosing establishes durable IgG and recallable MHC-I-specific CD8 memory.

To further understand the role of Arc in APC activation in response to tumor antigens, we analyzed donor DCs from WT, Arc-overexpressing, and Arc^−/−^ groups after stimulation with melanoma-derived EVs. As illustrated in Fig. 3A, primary DCs and MΦs were differentiated from bone marrow cultures supplemented with GM-CSF and IL-4, using cells from both wildtype (WT) and Arc^−/−^ knockout mice ^27^. Notably, Arc^−/−^ BM-DC/MΦs showed clear morphological differences compared to the WT cells (Fig. S7A). These donor cells were then divided into three experimental groups: (1) WT untransfected controls; (2) WT cells transfected with Arc and A5U for 16 h to induce VLV overproduction; and (3) Arc^−/−^ cells left untransfected, lacking Arc VLVs. All donor cell groups were subsequently exposed to melanoma antigens for processing and presentation. To obtain a broad spectrum of melanoma-associated antigens, B16-F10 culture supernatant was concentrated by ultrafiltration (100-kDa cutoff). This retains vesicles plus large secreted proteins/complexes (e.g., glycoproteins, HSP-peptide complexes), yielding a rich, heterogeneous antigen pool that is more reflective of the TME than EVs alone. To avoid confounding by carry-over melanoma EVs, we titrated the amount of B16-F10 supernatant added to DC2.4 and found that when the B16-F10 cell number exceeded 2× the DC2.4 cell number, transferring the concentrated supernatant introduced excess tumor EV carry-over. We therefore limited conditions ≤2× and processing steps that minimize residual B16 EVs in the DC-EV preparations, ensuring that downstream analyses reflect DC-derived vesicles (Fig. S8).

**Figure 3.**
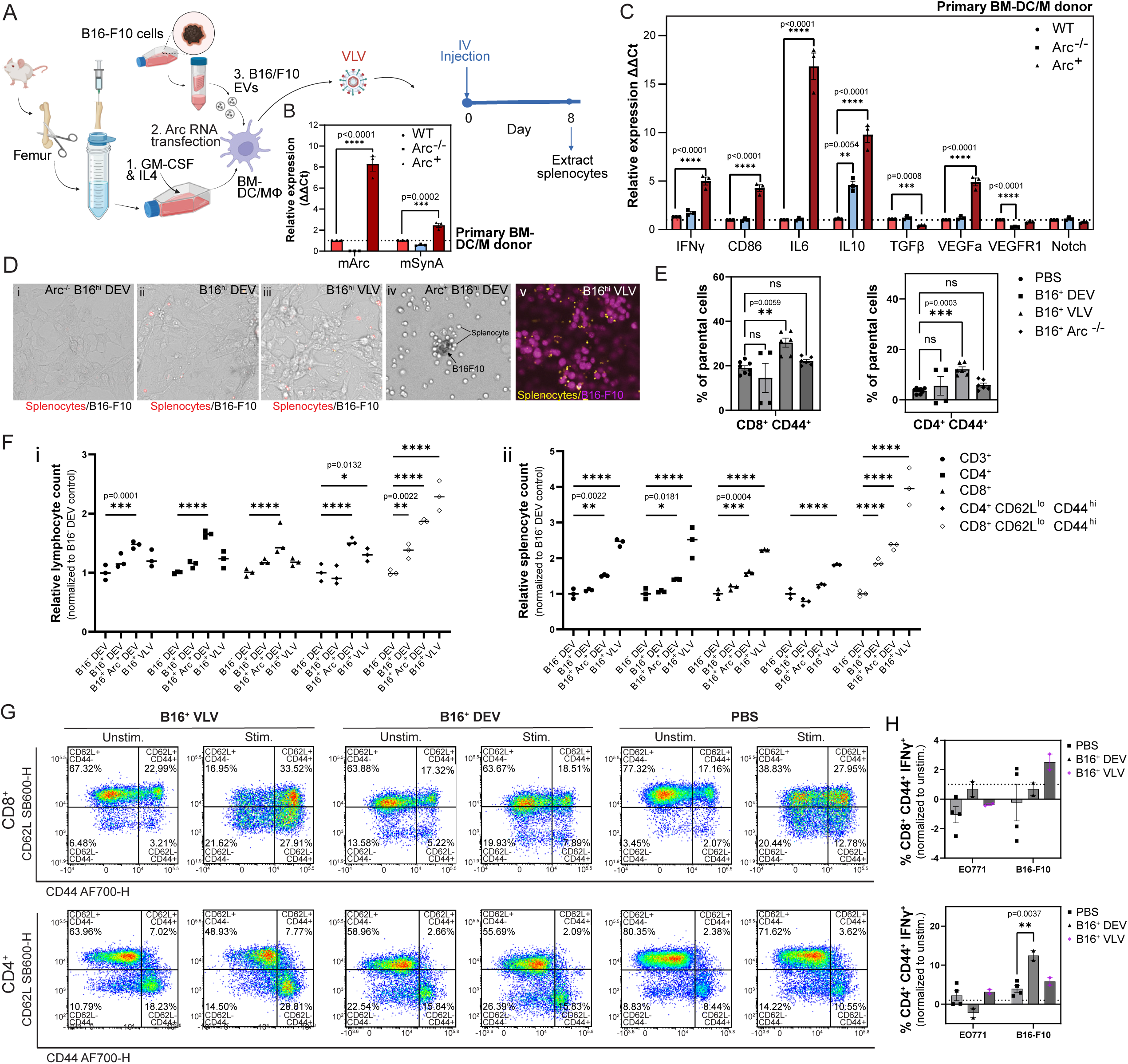
Arc is required for optimal immune activation and memory T cell generation. **(A)** Primary BM-DC/MΦs were differentiated from WT and Arc⁻/⁻ mice. WT cells were either transfected with *Arc* and *A5U-GFP* RNAs (Arc⁺ group) or left unmodified (WT group), whereas Arc-knockout cells remained unmodified as the Arc⁻/⁻ group. All groups were subsequently incubated with B16-F10 EVs/proteins to prepare vaccines for injection into WT recipient mice. **(B)** RT-qPCR analysis of mouse *Arc (mArc)* and *mSynA* expression in donor BM-DC/MΦs from the three groups, 24 h after exposure to B16-F10 EVs. Arc⁺ cells showed significantly elevated expression of both genes, whereas Arc⁻/⁻ cells showed reduced levels compared with WT controls. Mean ± SEM (*n* = 3). One-way ANOVA; p-values shown. **(C)** RT-qPCR of immune-related and signaling genes (CD86, IFN-γ, IL-6, IL-10, TGF-β, VEGFa, VEGFR1, Notch) in primary BM-DC/MΦs stimulated with B16-F10 EVs. Arc⁺ DCs exhibited strong upregulation of immune activation genes and reduced TGF-β, whereas Arc⁻/⁻ DCs showed comparable levels of most markers besides an IL-10 elevation and VEGFR1 reduction. Mean ± SEM (*n* = 3). One-way ANOVA; p-values shown. **(D)** Interaction assay between splenocytes from vaccinated mice and B16-F10 melanoma cells. CFSE-labeled splenocytes (red in i-iii; yellow in v) were co-incubated with NucSpot-labeled B16-F10 cells (purple in v). After 24 h, non-adherent cells were removed. WT and Arc⁺ splenocytes showed markedly increased tumor association, whereas Arc⁻/⁻ splenocytes did not. **(E)** Splenocytes isolated 8 days after melanoma vaccination were restimulated with B16-F10 EVs and analyzed by flow cytometry. Arc⁺ VLVs induced the strongest expansion of CD8⁺CD44⁺ and CD4⁺CD44⁺ T cells, whereas Arc^−/−^ DEVs did not trigger a detectable response. Mean ± SEM (*n* = 4-8). One-way ANOVA; p-values shown. **(F)** C57BL/6 mice received *i.v.* B16-F10-primed, DC2.4-derived DEVs or VLVs and were analyzed 72 h later in pooled lymph nodes (i) and the spleen (ii). Gating: singlets ◊ live ◊ CD45⁺ ◊ CD3⁺ ◊ CD4⁺/CD8⁺, with effector phenotype defined as CD62L^lo^ CD44^hi^. Arc⁺ VLVs significantly increased proliferation across CD3⁺, CD4⁺, CD8⁺, and effector subsets compared with B16⁻ DEV control. One-way ANOVA with Dunnett’s multiple comparisons; p-values shown. **(G)** Splenocytes isolated 12 days after vaccination were restimulated with B16-F10 EVs and analyzed for memory phenotypes. Representative plots show naïve (CD44⁻CD62L⁺), central memory (CD44⁺CD62L⁺), effector/effector memory (CD44⁺CD62L⁻), and pre-effector (CD44⁻CD62L⁻) subsets. Arc⁺ VLVs generated higher frequencies of CD8⁺ effector and central memory T cells. Data represents *n* = 4 mice per group. **(H)** Splenocytes isolated 12 days after vaccination were restimulated with B16-F10 or EO771 EVs. Arc⁺ VLVs induced significantly higher CD8⁺CD44⁺IFN-γ⁺ T-cell responses upon B16-F10 restimulation, with no response to EO771. CD4⁺CD44⁺IFN-γ⁺ cells expanded most strongly in the WT DEV group. Mean ± SEM (*n* = 2-4). One-way ANOVA; significant p-values shown.

After B16-F10 melanoma antigen exposure and an additional 24-hour culture in fresh serum-free medium, BM-DC/MΦs from each group were harvested for total RNA extraction and RT-qPCR analysis of immune-related genes and endogenous virus-like components, including the Arc capsid and the envelope protein Syncytin A (SynA). In Arc⁺ A5U⁺ transfected cells, mouse *Arc* (*mArc*) mRNA levels were substantially upregulated relative to WT non-transfected controls, and this increase was accompanied by elevated *mSynA* expression (Fig. 3B). In contrast, Arc⁻/⁻ BM-DC/MΦs showed no induction of *mArc* or *mSynA* following B16-F10 EV exposure. These data indicate that endogenous virus-like components in APCs are responsive to antigenic stimulation and that capsid and Env elements are functionally coupled.

At the same 24-hour timepoint, Arc⁺ BM-DC/MΦs displayed a broad transcriptional activation signature compared with WT controls, with increased CD86, IFN-γ, IL-6, IL-10, VEGFa, VEGFR1, and Notch, together with reduced TGF-β (Fig. 3C). This coordinated upregulation of co-stimulatory, inflammatory, and signaling genes, coupled to suppression of TGF-β, is consistent with an acutely activated DC state. By contrast, Arc⁻/⁻ BM-DC/MΦs expressed most activation markers to the same extent as WT DCs, indicating that Arc is not required for baseline DC activation. However, Arc deficiency altered the cytokine/signaling profile, with moderately elevated IL-10 and no reduction in TGF-β, suggesting a subtly dysregulated activation state. These findings support a model in which Arc is dispensable for initial DC activation but may be specifically required to establish the DC state that supports long-range, VLV-mediated T-cell engagement.

Accordingly, we next asked whether vesicles secreted by these DCs could trigger effector and memory T cell responses *in vivo*. Control DEVs and experimental VLVs generated from WT, Arc^+^ (Arc/A5U-overexpressing), and Arc⁻/⁻ BM-DC/MΦs were collected and systemically injected into WT C57BL/6J mice (6-8 weeks old, both genders). Eight days post-injection, splenocytes were isolated and co-cultured with B16-F10 melanoma cells. Both WT and Arc⁺ groups showed enhanced splenocyte-tumor interaction, whereas Arc⁻/⁻ did not, indicating Arc-dependent antigen-specific memory formation (Fig. 3D, S7B). For mechanistic analysis, splenocytes were restimulated *ex vivo* with B16-F10 EVs plus co-purified proteins for 6 h in serum-free OptiMEM, followed by 6 h in the presence of a protein transport inhibitor to permit intracellular cytokine accumulation. Live CD45⁺ splenocytes were gated and T cell subsets analyzed by surface activation/memory markers (CD3, CD4, CD8, CD44, CD62L) together with intracellular IFN-γ staining (Fig. S9A). At day 8, Arc⁺ VLV vaccination substantially increased CD4⁺CD44⁺CD62L⁻ and CD4⁺CD44⁺ populations relative to WT DEVs, whereas Arc⁻/⁻ DEVs reduced both CD4⁺CD44⁺ and CD8⁺CD44⁺ frequencies (Fig. S9B-C). However, day 8 proved too early to observe robust IFN-γ⁺ effector responses. We therefore shifted subsequent analyses to day 12 and optimized restimulation conditions by adding low-dose PHA-P (phytohemagglutinin-P, 0.5 µg/mL) to support T cell viability and extending antigen exposure to 12 h. Unstimulated splenocytes served as controls. By day 12, Arc⁺ VLV vaccination significantly expanded CD4⁺CD44⁺ and CD8⁺CD44⁺ populations compared with PBS controls, whereas Arc⁻/⁻ DEVs failed to induce increases (Fig. 3E). Having established Arc necessity using primary BM-DC/MΦ donors, we next assessed Arc sufficiency in gain-of-function assays using DC2.4 cells, selected for their ease of genetic manipulation and batch-to-batch consistency.

We first examined early *in vivo* responses to intravenously delivered, B16-F10-primed DC2.4-derived DEVs/VLVs, focusing on T-cell proliferation and early cytokine expression. 72 hours after *i.v.* injection into C57BL/6 mice, splenocytes and pooled cervical/axillary-brachial lymph nodes were analyzed. Flow cytometry revealed substantial and significant proliferation across multiple T cell subsets, including CD3⁺, CD4⁺, CD8⁺, and effector-phenotype CD4⁺/CD8⁺ (CD62L^lo^ CD44^hi^) populations, in the B16⁺ VLV group relative to controls (Fig. 3F i-ii). Consistent with early T cell priming after direct EV-T-cell engagement, RT-qPCR of pooled lymph nodes (six cervical and four axillary/brachial per mouse) showed a selective increase in *Il-2*, whereas *Il-4* trended upward without reaching significance and *Ifnγ* remained unchanged (Fig. S10). Together, these findings indicate that melanoma-primed VLVs rapidly expand multiple T cell subsets *in vivo* and induce an early IL-2 program, supporting their potential as fast-acting cancer vaccines.

To determine whether early expansion matured into antigen-specific memory, we restimulated day-14 splenocytes *ex vivo* with B16-F10 antigens after *i.v.* vaccination with DC2.4-derived, B16-F10-primed DEVs/VLVs. Following *ex vivo* restimulation, flow cytometric phenotyping showed increased frequencies of CD8⁺ CD44^hi^ CD62L^lo^ effector/effector-memory and CD8⁺ CD44^hi^ CD62L^hi^ central-memory T cells in the Arc⁺ VLV group compared with PBS and B16^+^ DEV controls (Fig. 3G). To further assess antigen specificity, splenocytes were restimulated with EO771-derived antigens in parallel with B16-F10 preparations. VLV vaccination induced a higher proportion of CD8⁺CD44⁺IFN-γ⁺ T cells upon B16-F10, but not EO771, restimulation, confirming antigen specificity (Fig. 3H). In contrast, B16⁺ WT DEVs elicited stronger CD4⁺CD44⁺IFN-γ⁺ responses than VLVs. This pattern is consistent with our MS data showing selective enrichment of Ova-derived peptides on MHC-I in VLVs, whereas Ova:MHC-II levels were comparable between VLVs and DEVs (Fig. 1E), indicating that Arc⁺ VLVs bias antigen presentation toward MHC-I pathways and preferentially enhance cytotoxic T cell immunity.

We carefully optimized the procedure for generating B16-F10-primed DEVs and VLVs for subsequent experiments, using DC2.4 cells for their ease of transfection, scalability, and reproducibility (Fig. 4A). We first performed EV flow cytometry to characterize how different B16-F10:DC2.4 cell ratios for priming (1×, 2×, and 5× B16-F10 cells per DC2.4 cell) affected the resulting vesicles. Because normalizing by total particle count would dilute DC-derived EVs in conditions with higher B16-F10 carry-over, we instead analyzed EVs collected from the same number of donor DC2.4 cells over the same culture period. Dye-only, no-stain EV controls and FMOs (fluorescence-minus-one controls) were included for gating. Across 1×, 2×, and 5× B16-F10:DC2.4 ratios, increasing B16-F10 input increased EV counts and right-shifted peaks for co-stimulatory molecules (CD80, CD86, CD40), and also increased MHC-II⁺ particle counts (Fig. 4B, S11). However, consistent with earlier findings, ≥2× B16-F10 input led to detectable B16-F10 EV carry-over in the final DEV preparations. We therefore proceeded with 1× (B16^lo^) and 2× (B16^hi^) conditions for subsequent experiments and used B16⁺ to denote the 1× condition when not otherwise specified.

**Figure 4.**
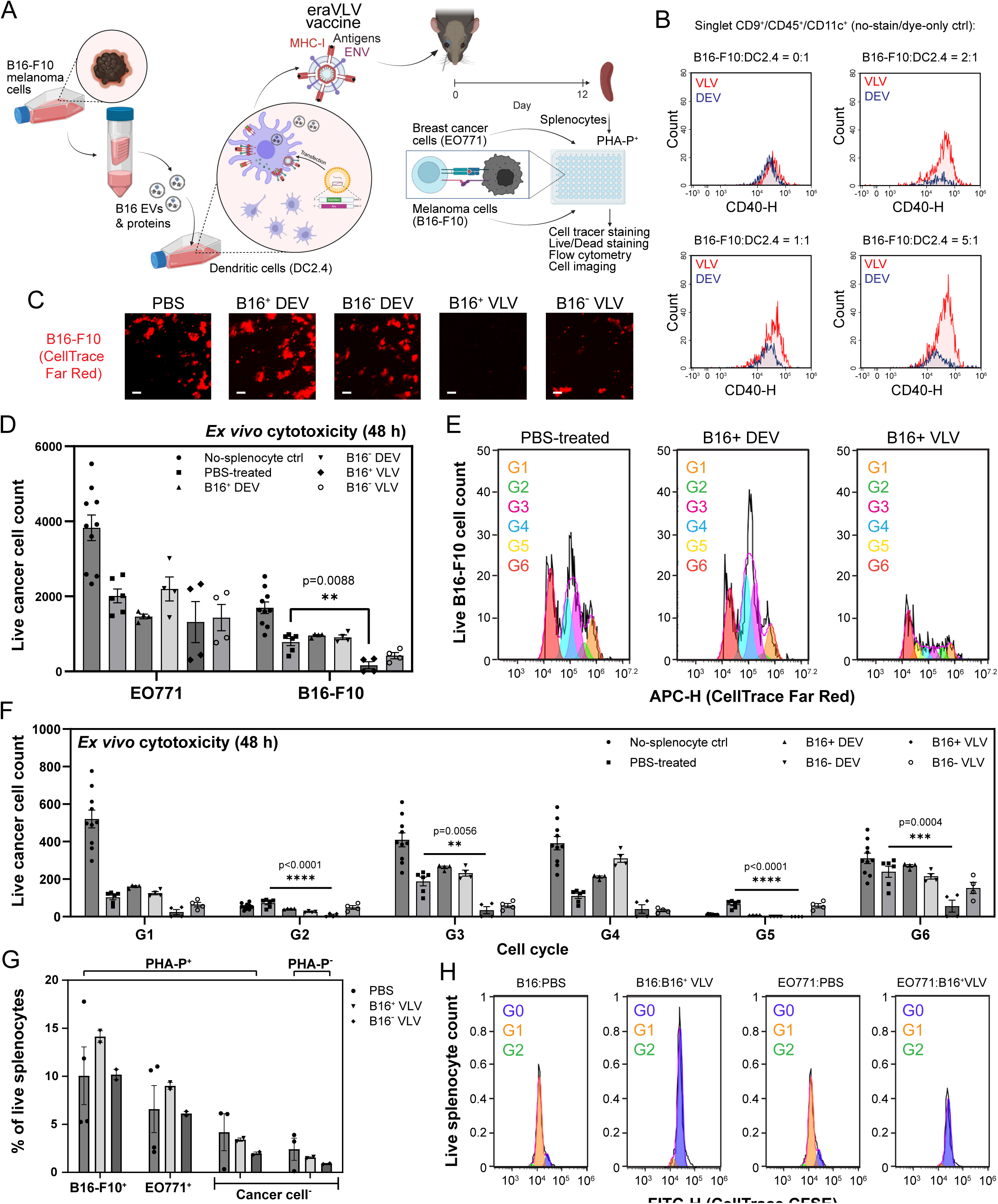
In vitro cytotoxicity of vaccine-induced splenocytes. **(A)** Schematic of the cytotoxicity assay. Splenocytes harvested on day 14 post-vaccination were labeled with CFSE and co-cultured with B16-F10 melanoma or EO771 breast cancer target cells pre-stained with CTFR at a total density of 0.6 × 10⁶ splenocytes per well in 100 µL (6 × 10⁶/mL), together with 3,000 tumor cells (bulk ratio ≈ 200:1). This high splenocyte density was selected to maintain *ex vivo* viability. Based on parallel phenotyping (≈ 3,000 IFN-γ⁺ CD8⁺ CD44⁺ cells per well), the effective effector-to-target ratio is ∼1:1 against 3,000 cancer cells. **(B)** EV flow cytometry of DC2.4-derived DEVs and VLVs generated at different B16-F10:DC2.4 cell ratios (0:1, 2:1, 1:1, 5:1). Histograms show CD40 signal on singlet CD9⁺CD45⁺CD11c⁺ vesicles (no-stain and dye-only controls indicated). Increasing the B16-F10:DC2.4 ratio shifts VLVs toward higher CD40 expression and increases CD40⁺ particle counts relative to DEVs. **(C)** Representative epifluorescence images of CTFR-labeled B16-F10 targets after 48 h co-culture with splenocytes from the indicated vaccine groups. Fewer red objects indicate greater tumor cell killing. Scale bars, 100 µm. **(D)** *Ex vivo* cytotoxicity at 48 h: live EO771 and B16-F10 target counts. Splenocytes from VLV-vaccinated mice significantly reduced B16-F10 cell numbers compared with PBS, B16⁺ DEV, or B16⁻ DEV controls, whereas EO771 targets showed no significant reductions. Symbols denote biological replicates (*n* = 4-10); bars show mean ± SEM. One-way ANOVA with Dunnett’s multiple-comparison test vs. control; exact P values are shown when significant. **(E)** CTFR dilution profiles of live B16-F10 targets after 48 h co-culture. Peaks G0-G6 denote successive division generations defined by dye dilution (proliferation generations). VLV-vaccinated splenocytes reduce surviving targets across generations, yielding lower counts, especially in early generations. **(F)** Generation-resolved quantification of live B16-F10 targets at 48 h based on CTFR dilution. Bars show surviving cells in each division generation (G1-G6) after co-culture with splenocytes from the indicated groups (no-splenocyte and PBS controls, B16⁺/B16⁻ DEVs, B16⁺/B16⁻ VLVs). Mean ± SEM; *n* = 4-10 biological replicates. Two-way ANOVA with Dunnett’s multiple comparisons vs. PBS (or no-splenocyte for baseline); significant p-values annotated. **(G)** Splenocyte viability in co-culture (% live CD45⁺ cells) with or without mitogenic support (PHA-P). VLV-vaccinated splenocytes maintain viability comparable to controls across target types. Mean ± SEM; *n* = 2-4. **(H)** CFSE dilution of splenocytes after 48 h co-culture showing minimal proliferation (G0-G2 peaks) under assay conditions, indicating that target-cell loss primarily reflects cytotoxicity rather than splenocyte expansion. Unless noted otherwise, all assays used 0.6 × 10⁶ splenocytes and 3,000 targets per well in 100 µL (bulk ratio ≈ 200:1; estimated effective E:T ≈ 1:1 for B16-F10).

To determine whether the immunologic skewing we observed translates into tumor cell killing, we co-cultured day-14 post-vaccination splenocytes with B16-F10 or EO771 target cells in the presence of PHA-P to maintain T cell viability (Fig. 4A). Representative CellTracker Far Red (CTFR) epifluorescence imaging at 48 h indicates a pronounced depletion of B16-F10 cells in the B16⁺ VLV condition (Fig. 4C), whereas PBS-splenocyte, B16⁺ DEV, and B16⁻ DEV wells exhibited similarly mild baseline loss (PHA-P^+^). Interestingly, B16⁻ VLVs produced a greater reduction in B16-F10 cells than either DEV control. Because the DC2.4 donor cells used to produce the EV vaccines are immortalized and partially transformed, they may express shared tumor-associated or stress antigens with B16-F10 melanoma, enabling cross-reactive cytotoxic responses that could account for the mild effect with B16⁻ VLVs (Fig. 4C). In addition to imaging, we performed flow cytometry. As expected, PHA-P induced modest baseline cytotoxicity in all groups. Against the PBS baseline, only splenocytes from B16⁺ VLV-vaccinated mice showed a significant, antigen-matched increase in B16-F10 killing at 48 h, whereas B16⁺ DEV or B16⁺ VLV splenocytes produced only a modest, non-significant effect against EO771 (Fig. 4D), consistent with specificity for the immunizing antigen. Using CTFR, we tracked target-cell divisions and observed an overall reduction across divisions with a disproportionate loss of early generations (G1-G5) and a relative skew toward G6 in the B16⁺ VLV group, indicating preferential elimination of early-dividing tumor cells, compared with the even generation profiles in PBS and B16⁺ DEV controls (Fig. 4E and S12A). Counts per generation were quantified (Fig. 4F), revealing that the B16⁺ VLV group led to significantly greater killing of B16-F10 cells across generations. Together, these show that Arc⁺ VLV vaccination converts immunologic bias into bona fide, antigen-specific tumor cell killing and proliferation arrest, with the strongest effects in the matched B16-F10 context.

On the responder side, within the live T cell gate (CFSE⁺), cancer^−^/PHA-P^−^ showed the lowest viability (expected *ex vivo*), whereas PHA-P alone increased viability, and adding tumor targets boosted it further, consistent with antigen-supported memory/effector cells); and B16⁺ VLV-primed splenocytes had the highest 48-h T cell viability, indicating the most resilient responder compartment, amplified by antigen encounter (Fig. 4G and S12B). On the other hand, T cell proliferation was attenuated (G0 retention) in both VLV treatment groups (EO771 and B16-F10) relative to PBS and DEV controls (Fig. 4H), suggesting an effector-skewed, non-expanding state under strong antigenic stimulation. Together with the selective tumor killing, this pattern supports a functional effector program that prioritizes cytotoxic activity over clonal expansion, consistent with rapid target engagement and sustained survival of antigen-experienced T cells in the B16⁺ VLV group.

Building on this potential, we next tested VLVs as *in vivo* cancer vaccines. A key advantage of this platform is its ability to adapt dynamically to each patient’s evolving tumor antigens, eliminating the need for predefined neoantigen targets. To assess efficacy, we used an unmodified orthotopic B16-F10 melanoma model. For vaccine production, donor DCs were exposed (serum-free) to B16-F10-conditioned supernatant containing both soluble tumor proteins and tumor-derived EVs, rich sources of diverse, clinically relevant tumor-associated antigens, and then processed as control versus Arc-engineered DCs to generate matched DEV and VLV formulations. We administered four doses of B16-F10⁺ DEV vaccines to C57BL/6 mice and, five days after the final dose, challenged them with B16-F10 melanoma cells to assess interventive (post-vaccination) efficacy (Fig. 5A), a preclinical analogue of treating or preventing tumor recurrence. To ensure unbiased data collection, tumor growth was quantified using double-blind caliper measurements independently conducted by multiple researchers, with further validation by micro-CT imaging that provides high-resolution confirmation of tumor size and morphology (Fig. 5A). Survival analysis revealed a substantial and significant advantage for mice treated with VLV vaccines, with a 60% survival rate compared to 37.5% for WT DEVs made with a high dose of B16-F10 EV antigens (B16^hi^ DEV, with a ratio of _donor_B16:_recipient_DC2.4 = 2:1) and 33.3% for B16^lo^ DEV control (B16:DC2.4 = 1:1) groups (Fig. 5B). In comparison, DEV⁻ control mice were euthanized between days 13-17 due to tumor volumes exceeding 1500 mm^3^. Comparative tumor size analysis showed a significant reduction in tumor growth in the Arc^+^ group, with daily measurements revealing a clear trend starting from five days post-B16 melanoma cell injection (Fig. 5C). By day 17, all DEV⁻ mice reached the experimental endpoint with tumors exceeding 1,500 mm³, confirming uncontrolled tumor progression in the absence of vesicle vaccination (Fig. 5D), whereas most sites in other sample groups remained tumor-free. For these groups, the largest tumor, in most cases the only tumor present, was extracted for size comparison and further analyzed through tissue sectioning and both histochemical and immunohistochemical (IHC) staining (Fig. 5E). Tumor volumes followed the order DEV⁻ > B16⁻ DEV > B16⁺ DEV > B16⁺ Arc⁺ DEV > B16⁺ Arc⁺ A5U⁺ VLV, indicating progressive enhancement of tumor control with VLV engineering. B16^hi^ Arc⁺ DEV tumors exhibited robust CD8⁺ T cell infiltration, whereas VLV-treated sites contained minimal residual tumor, largely skin adnexa (hair follicles) with only scattered CD8⁺ cells (Fig. 5E & G). This pattern likely reflects completion of the cytotoxic phase in the VLV group rather than reduced immunity, Arc⁺ DEVs represent an active antitumor phase, while VLVs achieve near-complete clearance, leaving few remaining targets. Interestingly, in the spleen of the same mice, CD4⁺ T cell numbers were comparable between Arc⁺ DEV and VLV groups (both exceeding controls), but CD8⁺ T cells were substantially lower in the VLV group (Fig. 5F & G). Such contraction of systemic CD8⁺ populations is consistent with post-clearance resolution, as antigen load and inflammatory cues decline once tumors are eliminated. Collectively, these data demonstrate that Arc-engineered B16^hi^ VLVs elicit stronger systemic and effector T cell responses, translating into superior tumor growth control.

**Figure 5.**
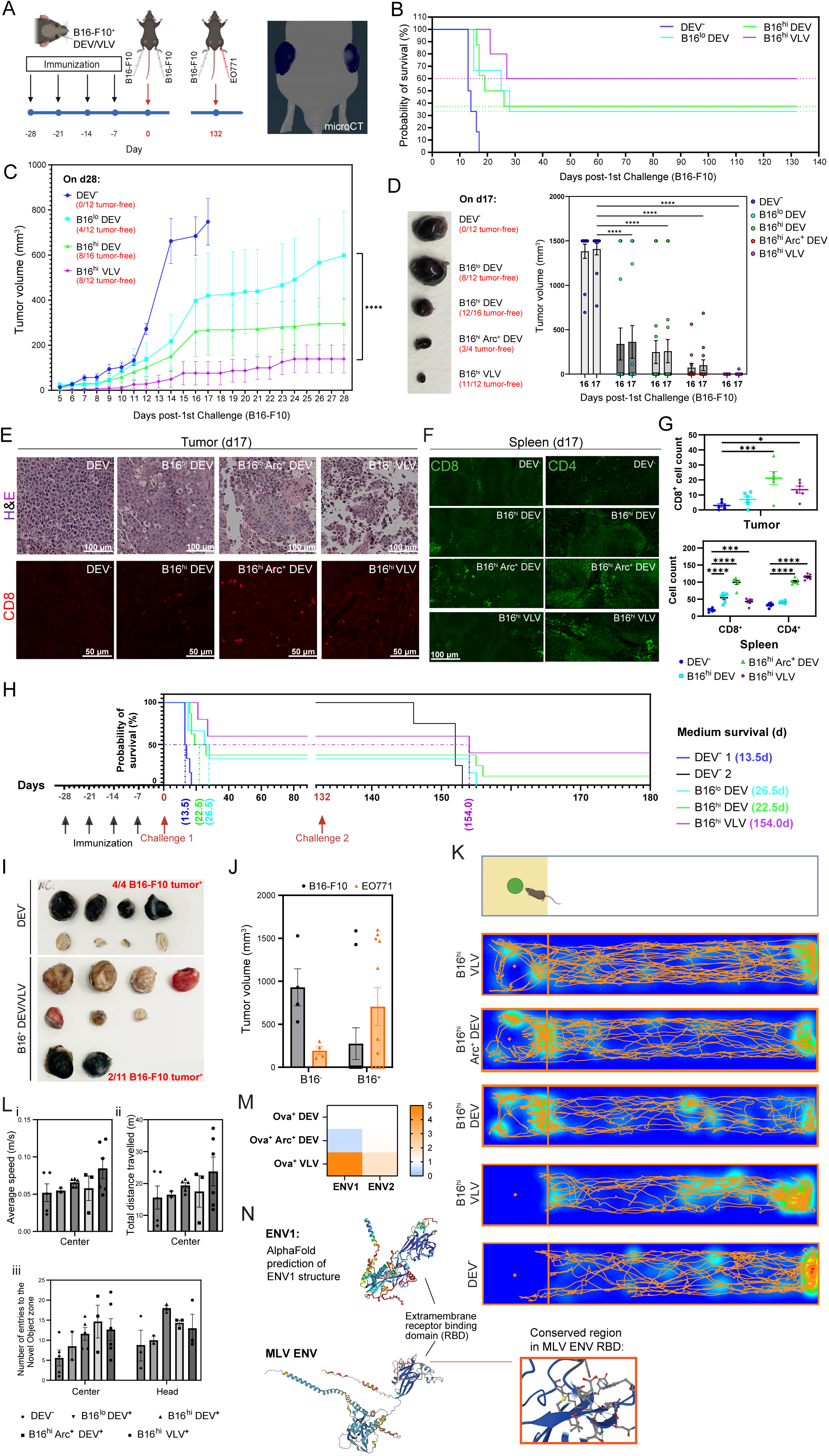
DC-derived VLV vaccines effectively prevent tumor growth. **(A)** shows the timeline and dosage schedule of B16-F10^+^ DEV vaccines administered to wildtype C57BL6 mice, alongside the initial injection of melanoma cells and rechallenge with both melanoma and breast cancer cells. Tumor measurements were conducted using micro-CT scanning and double-blind caliper measurements by multiple independent researchers to ensure data precision. **(B)** Kaplan-Meier survival curves were monitored to day 132 (pre-rechallenge). Mice receiving B16^hi^ VLVs showed a substantial survival advantage (75% injection sites tumor-free and 60% mice alive at day 132; median not reached) versus DEV⁻ NC (0%; median 13.5 d), B16^lo^ DEV (33.3%; median 26.5 d), and B16^hi^ DEV (37.5%; median 22.5 d). Pairwise log-rank (Mantel-Cox) tests *vs.* DEV⁻ NC: B16^hi^ VLV, **P = 0.0011; B16^hi^ DEV, **P = 0.0015; B16^lo^ DEV, *P = 0.0116. All DEV⁻ mice were euthanized on days 13-17 per IACUC tumor-size limits. Each mouse received two flank injections. N: DEV⁻ (6 mice/12 tumors), B16^lo^ DEV (6/12), B16^hi^ DEV (8/16), B16^hi^ VLV (6/12). 3 independent experiments were performed with some animals pre-assigned for imaging and staining were not followed for survival. Median survival (d): DEV⁻ (13.5) / B16^lo^ DEV (26.5) / B16^hi^ DEV (22.5) / B16^hi^ VLV (>study limit). **(C)** Tumor growth curves from day 5 post-implantation. Arc⁺ (B16^hi^ VLV) tumors remained significantly smaller than controls. Per-site measurements (two flanks per mouse); mean ± SEM. N (sites): 12, 12, 16, 12 for DEV⁻, B16^lo^ DEV⁺, B16^hi^ DEV⁺, B16^hi^ VLV⁺, respectively (3 independent experiments). Statistics: two-way mixed-effects ANOVA (Dunnett’s multiple-comparisons); B16^lo^ DEV *vs.* B16^hi^ VLV: ****P < 0.0001. **(D)** When the final DEV⁻ control mouse reached humane endpoint on day 17, tumors were harvested for cross-sectional comparison. **Left**: representative tumors from one mouse per group (chosen a priori as the animal bearing the only or largest tumor for photographic comparison). **Right:** quantification of all injection sites at day 17 shows DEV⁻ > B16^lo^ DEV > B16^hi^ DEV > B16^hi^ Arc⁺ DEV > B16^hi^ VLV (mean ± SEM). Measurements were by caliper volume 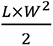 at the site level. n (sites): DEV⁻ = 12; B16^lo^ DEV⁺ = 12; B16^hi^ DEV⁺ = 16; B16^hi^ Arc⁺ DEV⁺ = 12; B16^hi^ VLV⁺ = 12. Statistics: two-way mixed-effects ANOVA with Dunnett’s multiple-comparisons test. \*\*\*\**P*< 0.0001. **(E)** H&E: Arc⁺ VLV tumors were smaller with reduced viable tumor area, lower tumor-cell density, and less cohesive architecture compared with controls. CD8 IHC: A significant increase in CD8⁺ T cells was detected in B16^hi^ Arc⁺ DEV and VLV tumors *vs.* DEV⁻ controls. Notably, section-wide CD8⁺ counts in the VLV group appeared lower than Arc⁺ DEV, reflecting the scarcity of residual viable tumor in VLV sections (large areas comprised skin adnexa such as hair follicles and connective stroma). **(F)** Spleens were collected from the same tumor-bearing mice for IHC staining of CD4⁺ and CD8⁺ T cells. Splenic CD8⁺ T cell counts followed the order DEV⁻ < B16^lo^ DEV < B16^hi^ DEV = B16^hi^ Arc⁺ DEV, while the B16^hi^ Arc⁺ A5U⁺ VLV group displayed minimal CD8⁺ T cell staining. CD4⁺ T cells followed the order DEV⁻ < B16^lo^ DEV < B16^hi^ DEV < B16^hi^ Arc⁺ DEV = B16^hi^ Arc⁺ A5U⁺ VLV. **(G)** CD8⁺ cells in 5 representative tumor regions were quantified from CD8 IHC. Splenic CD4⁺ and CD8⁺ cells were counted in 5 representative regions. Mean ± SEM. Statistics: one-way ANOVA with Dunnett’s multiple-comparison, exact *P* values in panel. **(H)** Survival following challenge with melanoma cells and rechallenge with both melanoma and breast cancer cells: (i) In one of three independent experiments, long-term survivor mice were rechallenged with B16-F10 melanoma and EO771 breast cancer cells on day 132. A second batch of DEV⁻ controls was introduced as members of the first batch succumbed. (ii) Median survival times were as follows: 13.5 days for the first batch of DEV⁻ control (NC1); 14 days for the second batch of DEV⁻ control (NC2); 26.5 days for B16^lo^ DEV control; 22.5 days for B16^hi^ DEV control; and 154 days for the B16^hi^ VLV group. **(I)** Representative tumors collected at euthanasia from control and vaccinated mice. Each image shows all excised melanoma (B16-F10) and breast cancer (EO771) tumors from one representative experiment, arranged by group (DEV⁻ control *vs.* B16 ^hi^ DEV/VLV-vaccinated). **(J)** Quantification of tumor volumes at euthanasia. Control mice (DEV⁻) showed rapid melanoma progression and developed large tumors, leading to early mortality. In contrast, vaccinated groups (B16^hi^ DEV or B16 ^hi^ VLV) showed strong protection against melanoma, substantially extending the medium survival. This prolonged survival allowed the slower-growing breast cancer (EO771) tumors to reach larger sizes in some mice. **(K)** In addition to monitoring tumor growth, the behavioral well-being of treated mice was assessed, including locomotion and cognitive capacities using a novel object test. The movement tracks (orange) provide a visual representation of the paths taken by the mice within the test chamber. The heat map shows the density of movement within different areas, indicating the regions where the mice spent the most time. Besides the negative control mice, which are from new cancer cell injections, subjects in all experimental groups studied here are treated survivors after two rounds of tumor rechallenge. Representative tracks are shown here, with additional tracks available in the supplementary figure. **(L)** B16^hi^ vaccinated survivor mice exhibited increased average speed (i), total travel distance (ii), and number of entries into the novel object zone (iii), compared to control mice. The VLV group exhibited the highest level of locomotion activity. N = 3-5 in each sample group. Mean ± SEM. **(M)** MS label-free quantification enabled the identification and quantification of EV membrane proteins. The heat map provides a comparative analysis of protein expression levels across different sample groups. The heat map shows an upregulation of endogenous Envs in Ova-stimulated VLVs, compared to both the Ova^‒^ DEV control and the Arc^+^/A5U^‒^ DEV control. **(N)** Structural comparison of the MLV envelope (bottom) and AlphaFold prediction of ENV1 (top). MS analysis revealed a high degree of sequence similarity in the extracellular receptor-binding domain between MLV Env and a specific sequence enriched in Ova^+^ VLVs (ENV1). A beta-strand sequence within this domain was identified as 100% conserved, highlighting significant structural conservation critical for receptor interaction.

At day 132 after the initial B16-F10 challenge, no PBS controls remained tumor-free (0/6 mice), whereas B16^lo^ DEV protected 2/6 mice (33.3%; 4/12 sites), B16^hi^ DEV protected 3/8 mice (37.5%; 7/16 sites), and B16^hi^ VLV protected 4/6 mice (66.7%; 9/12 sites). To assess durability and breadth of protection, surviving mice were rechallenged: each animal received B16-F10 intradermally on one flank and EO771 subcutaneously on the contralateral flank. After the original cohort had succumbed, we introduced a second, contemporaneous DEV^‒^negative control cohort to anchor rechallenge readouts to matched controls. As previously demonstrated, B16^hi^ VLVs selectively protected against melanoma, with minimal cytotoxicity toward EO771 cells *in vitro* (Fig. 4D).

Interestingly, *in vivo* VLV vaccination prevented progression of both melanoma and breast cancer. It extended median survival up to 11-fold *vs.* the DEV^‒^ control and nearly 7-fold versus the B16^hi^ DEV control, with 67% of injection sites (33% of mice) remained tumor-free after a second dual-tumor rechallenge (Fig. 5H). Exclusively *in vivo*, this cross-tumor protection most likely reflects bystander effects arising from the inflammatory and antigen-presenting context created by VLVs. First, Arc⁺ VLVs potently activate and license DCs, increasing co-stimulation and cytokines (e.g., type I IFN, IL-12), which can drive epitope spreading: Once the melanoma lesion regressed, dying cells provided antigen for cross-priming and epitope spreading to shared self-peptides and cross-reactive motifs, broadening CD8⁺ responses beyond melanoma and enabled control of the breast tumors. Second, VLVs upregulate adhesion molecules and leukocyte-interacting proteins on vesicles and host cells, enhancing T cell trafficking and retention in distant lesions; the same pro-inflammatory milieu (IFN-γ/TNF) can impose bystander cytotoxicity on neighboring tumor cells regardless of their cognate antigen. Third, innate arms likely contribute: VLVs can trigger NK-cell activation and macrophage reprogramming, amplifying antigen-independent killing and phagocytosis across tumor types. Finally, the presence of shared tumor antigens between melanoma and breast cancer may further contribute to the protective efficacy of our VLV vaccine across multiple cancer types, supporting its potential as a broad-spectrum therapy ^28–34^. Nonetheless, we observed preferential protection against melanoma overall. In the PBS-treated control group, mice rapidly reached euthanasia criteria due to aggressive melanoma tumor growth exceeding the permissible size limit, whereas the less aggressive breast cancer tumors remained small (Fig. 5I-J). However, vaccination with B16-F10^+^ DEVs and VLVs showed substantial protection against melanoma, extending the survival periods of treated mice significantly (Fig. 5H), allowing them more time to grow larger breast cancer tumors. Eventually, these vaccinated mice reached euthanasia criteria with significantly larger breast cancer tumors (Fig. 5I-J). Collectively, these results establish VLVs as a potent vesicle vaccine that triggers durable, cross-tumor immunity, substantially prolonging survival and yielding tumor-free survivors even after dual-tumor rechallenge. Such antigen-agnostic, polyclonal responses may be particularly valuable for tumors with high mutational burden or antigenic drift, where immune escape through neoantigen evolution limits the durability of conventional single-antigen vaccines.

In addition to monitoring tumor growth in vaccinated, tumor-bearing mice, we assessed their locomotive and cognitive wellness using a novel object behavior test. Mice treated with B16^hi^ DEV vaccines exhibited enhanced exploratory behavior in the novel object behavior test, displaying higher frequencies of interaction with the novel object (Fig. 5K & S13). They showed increased speed (Fig. 5L i), travel distance (Fig. 5L ii), and number of entries into the novel object zone (Fig. 5L iii). In contrast, mice in the DEV^‒^ control group tended to avoid the novel object zone. Altogether, these suggest that VLVs not only inhibit tumor growth but also exert positive effects on cognitive function and overall well-being in treated mice.

In our further exploration of the molecular mechanisms underlying the formation and immune competence of VLV vaccines, we found that combined antigen stimulation and Arc overexpression reactivated endogenous Envs, which became incorporated into VLV membranes and may engage T cell receptors in a manner reminiscent of classical retroviral Envs. This observation is consistent with prior findings that expression of endogenous retroviral elements (ERVs) is frequently reactivated by immune stimulation or by relaxation of epigenetic silencing, leading to renewed transcription of virus-like genes and double-stranded RNA intermediates ^35,36^. Membrane proteins were enriched from Ova-primed DC-derived vesicles using the Pierce™ Mem-PER™ kit, which was originally designed for cellular membranes. Because vesicle membranes cannot be pelleted by low-speed centrifugation, the protocol was adapted by incorporating an ultrafiltration step to collect membrane fragments. This modification, however, prevents the complete removal of high-molecular-weight intra-vesicular proteins, so the resulting preparation represents a membrane-enriched rather than strictly membrane-exclusive fraction. Subsequently, Label-free MS identified peptide fragments with high homology to the extracellular receptor-binding domain of murine leukemia virus (MLV) Env, specifically enriched in Ova⁺ VLVs (Fig. 5M; ENV1, annotated as “MLV-related proviral Env polyprotein-like”). Notably, a beta-sheet motif within this domain was 100% conserved (Fig. 5N). In addition to this sequence, an extracellular peptide sequence from the endogenous porcine Env was also increased in VLVs (Fig. 5M, ENV2, or putative envelope polyprotein). The natural targets of MLV include B cells and T cells ^37,38^. The conservation between retroviral Envs and Arc-associated DEV membrane proteins suggests that VLV vaccines may be exploiting similar pathways used by retroviruses to engage and activate immune cells. Importantly, the upregulation of Envs was observed exclusively in VLVs when both the Arc capsid protein and Arc 5’ UTR were co-introduced, suggesting that the 5’ UTR of Arc plays crucial roles in Env recruitment. Moreover, the upregulation of several membrane proteins involved in leukocyte adhesion and immune surveillance in Arc^+^ DEVs suggests complex interactions between VLV components and immune activation pathways. Such molecular mimicry opens new avenues for understanding how DEV vaccines can be fine-tuned to harness virus-like properties for improved vaccine efficacy against diseases like cancer.

We profiled melanoma antigen-presenting vesicles by MS. Proteins detected in the melanoma dataset were cross-referenced with our Ova dataset and the ExoCarta, Vesiclepedia, and EVpedia repositories, revealing both canonical EV components and newly observed vesicular proteins (Fig. 6A and S14). Differentially abundant proteins (UniProt accessions) from each pairwise comparison (B16^−^ DEV *vs.* B16^+^ DEV; B16^+^ DEV *vs.* B16^+^ VLV; B16^−^ DEV *vs.* B16^+^ VLV) were compiled and submitted to G:Profiler (g:GOSt) for GO:BP enrichment against the full quantified proteome background. Multiple testing was controlled (FDR), and terms with adjusted *P* < 0.05 were reported (Fig. 6C; Table S3). As shown in Fig. 6C(i), enrichment of specific biological processes in B16⁺ VLVs compared to B16⁻ DEVs, such as *response to virus* and *cytoplasmic pattern-recognition receptor signaling pathways*, indicates that VLVs leverage viral mimicry to enhance innate immune detection and response. In B16⁺ VLVs, enrichment of *organonitrogen compound metabolism* likely reflects heightened amino-acid/peptide metabolic programs that fuel antigen processing and proteostasis, processes tightly coupled to MHC-I peptide generation in APCs. Concurrent enrichment of *non-membrane-bounded organelle assembly* is consistent with increased formation of ribosome/stress-granule-like RNP condensates that support translational control and proteome remodeling during vesicle biogenesis and antigen handling. As shown in Fig. 6C(ii), the most significantly enriched GO:BP term in the B16⁺ VLV *vs.* B16⁺ DEV comparison was *response to virus*. This enrichment likely reflects the intrinsic antiviral transcriptional program activated by the overexpression of VLV components within VLV-producing DCs. Enrichment of *positive regulation of gene expression* and *TNF production* aligns with NF-κB/IRF pathway activation commonly observed after viral-mimic stimulation (e.g., poly I:C), which rapidly induces TNF-α and downstream inflammatory cues that recruit and activate immune cells. Terms such as *peptide metabolism*, *ribosome assembly*, and *protein localization* are compatible with enhanced antigen-processing capacity: ongoing translation supplies defective ribosomal products (DRiPs) and other substrates for the (immuno)proteasome-MHC-I pathway, while boosted trafficking supports peptide loading and export. Taken together, the VLV condition behaves like an adjuvanted, virus-mimicking vesicle, not merely presenting tumor antigens but also triggering innate antiviral circuitry that amplifies adaptive priming. As shown in Fig. 6C(iii), B16⁺ DEVs relative to B16⁻ DEVs showed enrichment of pathways such as *blood coagulation*, *TGF-β signaling regulation*, and *MAPK cascade modulation*, indicating that the addition of melanoma antigens enhances vesicle functions related to cell signaling, migration, and intercellular communication. Enrichment of *metal-ion response* and *acute-phase reaction* terms is consistent with DC activation by tumor-associated antigens, which often triggers redox- and cytokine-regulated stress responses. Collectively, these findings suggest that melanoma antigen incorporation primes DC-derived VLVs for efficient antigen presentation and immunogenic communication with T cells, while the addition of virus-like components further amplifies innate antiviral and inflammatory signaling, enhancing immune cell recruitment and activation beyond basal presentation capacity.

**Figure 6.**
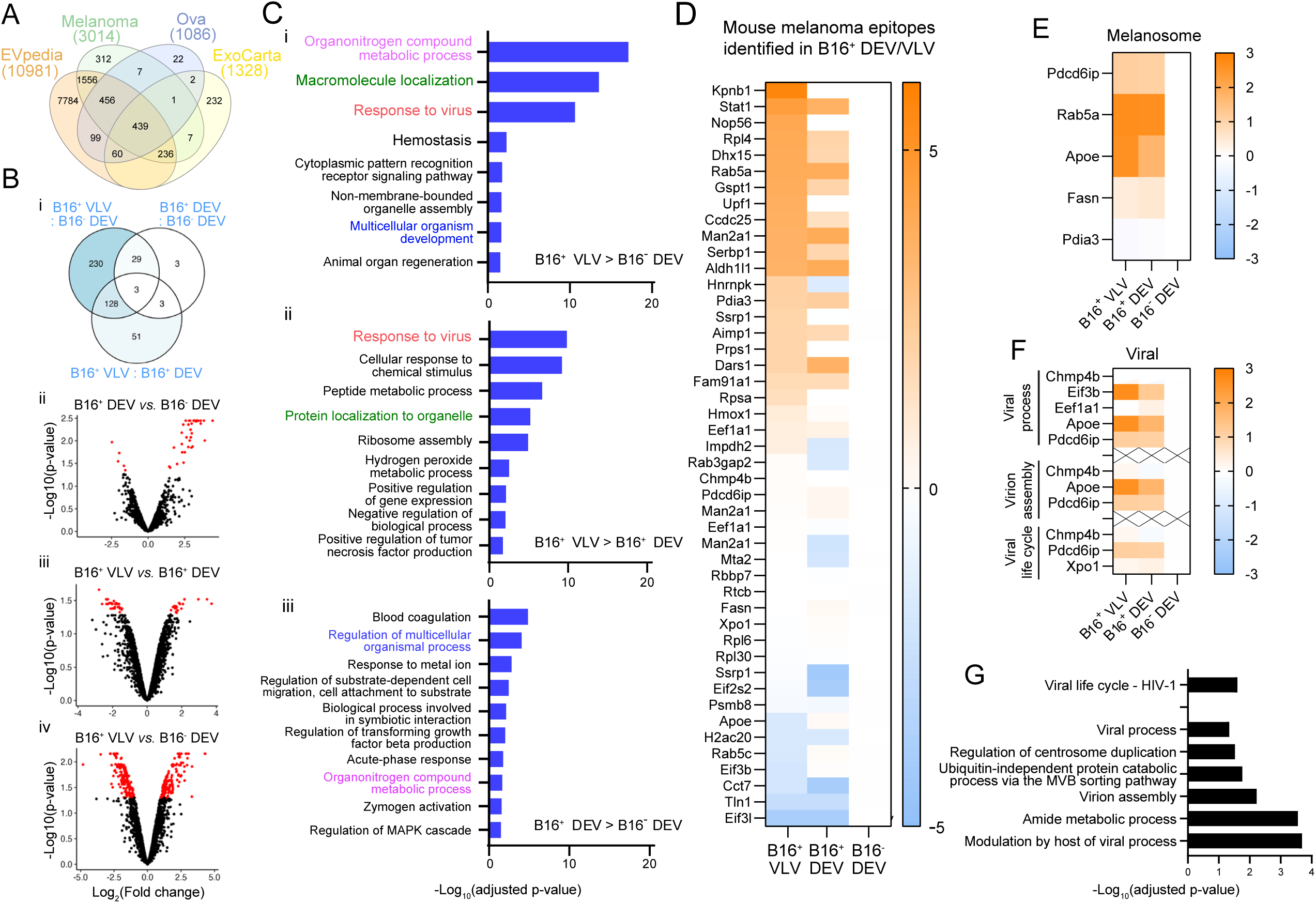
Proteomic and immunogenic profiling of melanoma antigen-containing vesicles. **(A)** Venn diagram compares the proteins identified in the B16^+^ and Ova^+^ DC-derived vesicles with those annotated in the ExoCarta and EVpedia databases. **(B)** Differential proteomics. **(i)** Venn diagram of proteins meeting the significance criteria across pairwise contrasts (B16⁺ VLV *vs.* B16⁺ DEV; B16⁺ DEV *vs.* B16⁻ DEV; B16⁺ VLV *vs.* B16⁻ DEV). **(ii-iv)** Volcano plots for **(ii)** B16⁺ DEV *vs.* B16⁻ DEV, **(iii)** B16⁺ VLV *vs.* B16⁺ DEV, and **(iv)** B16⁺ VLV *vs.* B16⁻ DEV. Axes show log₂(fold change) and −log₁₀(adjusted p). Proteins exceeding a 1.5-fold cutoff with FDR < 0.05 (Benjamini-Hochberg correction) are highlighted in red. Data represent two independent vesicle preparations per group analyzed by LC-MS/MS under identical conditions. **(C)** GO:BP enrichment analysis of differentially expressed vesicle proteins. Bars indicate −log₁₀ (adjusted p) values for significantly over-represented processes. **(i)** Processes enriched in B16⁺ VLVs *vs.* B16⁻ DEVs, highlighting viral-response, macromolecule-localization, and multicellular-development pathways. **(ii)** Processes enriched in B16⁺ VLVs *vs.* B16⁺ DEVs, including protein localization to organelles, ribosome assembly, and TNF-α production regulation. **(iii)** Processes enriched in B16⁺ DEVs *vs.* B16⁻ DEVs, featuring coagulation, MAPK-cascade regulation, and metabolic responses. Only terms with adjusted p < 0.05 are shown. **(D)** Heat map of normalized log₁₀ (fold change) values for 46 melanoma epitopes identified in B16⁺ DC2.4 vesicles and matched to the IEDB mouse melanoma antigen library. Many epitopes display stronger representation in B16⁺ VLVs compared with DEVs, consistent with enhanced antigen packaging. **(E)** GO: Cellular Component analysis of parent proteins from identified epitopes indicates strong enrichment for the *melanosome* category in both B16⁺ VLVs and DEVs, reflecting selective loading of melanoma-associated cargoes (e.g., Pdia3, Rab5a, Apoe, Fasn). **(F)** Functional clustering of vesicle proteins shows preferential enrichment of viral-process pathways in B16⁺ VLVs, including virion assembly, MVB-sorting, and viral-life-cycle-related functions (e.g., Chmp4b, Pdcd6ip, Eif3b, Xpo1). These categories support a viral-mimicry signature consistent with endogenous retroelement-derived vesicle formation. **(G)** Additional enriched biological processes (false-discovery-adjusted p < 0.05) include modulation by host of viral process, ubiquitin-independent protein catabolism, and centrosome-duplication regulation. Bars represent −log₁₀(adjusted p), truncated at 5 for display clarity.

Focusing on specific targets, comparative proteomic analysis between B16⁺ and B16⁻ DEVs revealed elevated expression of endocytosis- and trafficking-related proteins, including *Igf2r*, *Lcat*, *Fgb*, *Lum*, *Thbs1*, and *Fn1*, indicating enhanced uptake and processing of melanoma-derived antigens (Table S3). Upregulated proteins associated with proteolysis, peptidase activity, and cross-presentation of exogenous antigens, such as *Ctnnb1*, *Psmb1*, *Psmb2*, *Psmb10*, *Klkb1*, *F2*, *Hgfac*, *Gsk3a*, *Ufd1*, *Serpinf1*, *Gsn*, *Mug1*, *Thbs1*, *Itih2*, *Fn1*, *Fbln1*, *Serpinc1*, *Pzp*, and *Itih4*, further support enhanced antigen processing and presentation capacity in melanoma antigen-loaded DEVs. The concurrent enrichment of immune activators (*C1qtnf3*, *Serpinc1*) and metabolic regulators (*Mat1a*, *Aldob*) suggests coordinated adaptation of immune and metabolic pathways to sustain antigen handling and vesicle maturation. By contrast, VLVs were enriched for antiviral and interferon-stimulated proteins, *Gbp2*, *Iigp1*, *Ifit1*, consistent with innate antiviral programs driven by the virus-like payload. We also saw higher *Stat1/2/3*, *Eif2ak2* (PKR), *Trim25*, and *Oasl1*, marking activation of interferon and cytokine-mediated signaling pathways and indicating broad immune amplification. Increased cytoprotective factors such as *Hmox1* and *Gclm* point to oxidative stress adaptation during immune activation. Together with *Sqstm1*, *Serpinc1*, and *Serpinf1*, these changes indicate stress- and inflammation-regulatory circuits that facilitate cellular homeostasis while sustaining immune surveillance.

MS-identified peptides in B16⁺ DEVs and VLVs were queried against the Immune Epitope Database (IEDB) for mouse melanoma epitopes: several peptides from B16⁺ vesicles matched established immunogenic sequences, supporting their capacity to trigger melanoma-specific responses (Fig. 6D). Across conditions we detected hundreds of shared proteins, with 16 significantly increased in B16⁺ VLVs versus B16⁻ DEVs (Fig. 6D). GO analysis of these proteins and the shared epitopes highlighted compartments and processes relevant to melanoma and VLV biology. Notably, *melanosome* was strongly enriched, with melanosome-associated proteins increased in B16⁺ VLVs (Fig. 6E), consistent with improved display of melanoma antigens. The enrichment of GO terms such as *modulation by host of viral process*, *virion assembly*, and *viral life cycle (HIV-1)* indicates that VLV proteins engage pathways commonly hijacked or mimicked by viruses to assemble and release particles (Fig. 6F). The presence of *ubiquitin-independent protein catabolic process via the multivesicular body (MVB) sorting pathway* points to enhanced ESCRT-mediated vesicle formation, a mechanism shared by both exosomes and many enveloped viruses. *Amide metabolic process* suggests elevated protein and peptide synthesis supporting antigen processing, while *regulation of centrosome duplication* reflects cytoskeletal remodeling linked to vesicle trafficking and secretion (Fig. 6G). Collectively, these enrichments support a viral-mimicry program within VLVs that promotes vesicle biogenesis, antigen presentation, and immune activation.

## Discussion

Our study uncovers a previously unrecognized mechanism by which DCs repurpose ancient viral components into endogenous VLVs to amplify adaptive immunity. We show that Arc^+^ VLVs function as endogenous mediators of long-range antigen presentation, directly engaging T cells. Through comparative gain-and loss-of-function analyses, we demonstrate that Arc is essential for this process, Arc^‒/‒^ knockout DEVs failed to generate antigen-specific T cell memory *in vivo*, whereas overexpression of Arc and A5U markedly increased the proportion of Arc⁺ VLVs among total DEVs (>70%). These engineered VLVs confer multiple immunologic advantages: enhanced MHC-I antigen presentation, robust immune-pathway activation and durable humoral immunity (**∼**257-fold increase over control by week 13) and sustained memory formation. In melanoma models, Arc⁺ VLV vaccination produced antigen-specific cytotoxicity *ex vivo*, near-complete tumor regression *in vivo* in 75% of injection sites after the initial melanoma challenge and an up to 11-fold extension of survival, with 67% of injection sites tumor-free after a second dual-tumor (melanoma and breast cancer) rechallenge. Notably, melanoma-primed VLVs also provided bystander protection against breast cancer *in vivo*, demonstrating cross-tumor efficacy and adaptability to heterogeneous or mutating antigenic landscapes. By recruiting endogenous envelope proteins and activating interferon-driven antigen-processing networks, VLVs deliver intrinsic innate co-stimulation that amplifies adaptive priming without exogenous adjuvants. Collectively, these results establish Arc VLVs as a powerful, antigen-agnostic and self-adjuvanting vesicle vaccine platform with broad translational potential for durable cancer immunotherapy.

## Limitations of the study

This study has limitations. First, although membrane-enriched mass spectrometry identified Env-like proteins (ENV1/ENV2) selectively increased in Arc⁺ VLVs, we did not directly test whether these Envs mediate T cell engagement. Blocking experiments using antibodies or soluble receptor ectodomains for candidate receptors will confirm binding mechanisms. Second, although Arc rescue experiments were performed using primary Arc⁻/⁻ BM-DC/MΦs, neither WT nor Arc-rescued Arc⁻/⁻ DEVs produced strong splenic enrichment or melanoma-specific memory. We therefore did not pursue rescue experiments for the interventive melanoma vaccine, especially considering the seven-day differentiation required for primary BM-DC/MΦs made them impractical for weekly vaccine production.

## STAR★METHODS

### Key resource table

| REAGENT or RESOURCE | SOURCE | IDENTIFIER |
| --- | --- | --- |
| Antibodies (flow cytometry) |  |  |
| Anti-mouse CD40, SB436, clone 1C10 | Thermo Fisher Scientific<br>(Invitrogen/eBioscience) | 62-0401-82 |
| Anti-mouse CD80, PE, clone 16-10A1 | Thermo Fisher Scientific | 12-0801-82 |
| Anti-mouse CD86, APC, clone GL-1 | Thermo Fisher Scientific | 17-0862-82 |
| Anti-mouse CD45, AF700, clone 30-F11 | Thermo Fisher Scientific | 56-0451-82 |
| Anti-mouse CD11c, BV650, clone N418 | Thermo Fisher Scientific | 416-0114-80 |
| Anti-mouse CD11b, AF488, clone M1/70 | Thermo Fisher Scientific | 53-0112-82 |
| Anti-mouse F4/80, Super Bright 600, clone BM8 | Thermo Fisher Scientific | 63-4801-82 |
| Anti-mouse MHC-II (I-A/I-E), APC-eFluor 780, clone M5/114.15.2 | Thermo Fisher Scientific | 47-5321-82 |
| Anti-mouse MHC-I (H-2K<sup>b</sup>), PE-Cy7, clone AF6-88.5.5.3 | Thermo Fisher Scientific | 56-5958-82 (*AF6-88.5.5.3*) |
| Anti-H-2K<sup>b</sup>:SIINFEKL (OVA:MHC-I complex), PE, clone 25-D1.16 | Thermo Fisher Scientific (eBioscience) | 12-5743-82 |
| Anti-mouse CD115 (CSF1R), PE-Cy7, clone AFS98 | Thermo Fisher Scientific | 25-1152-80 |
| **Viability/compensation and buffers (flow)** |  |  |
| LIVE/DEAD™ Fixable Lime Dead Cell Stain | Thermo Fisher Scientific | L34989 |
| UltraComp eBeads™ Compensation Beads | Thermo Fisher Scientific | 01-3333-42 |
| ArC™ Amine Reactive Compensation Bead Kit | Thermo Fisher Scientific | A10628 |
| FcR Blocking Reagent (mouse) | Miltenyi Biotec | 130-092-575 |
| Flow Staining Buffer | Thermo Fisher Scientific | 00-4222-57 |
| Complete Staining Buffer (SB) | Thermo Fisher Scientific | SB-4401-42 |
| **Chemicals, peptides, and recombinant proteins** |  |  |
| Lipopolysaccharide (LPS) | Sigma-Aldrich | L5543 |
| Ovalbumin (OVA) | InvivoGen | vac-stova, vac-sin, vac-isq |
| PHA-P (phytohemagglutinin-P) | InvivoGen | inh-phap-2 |
| β-mercaptoethanol | Gibco | 21985023 |
| **Critical commercial assays & kits** |  |  |
| HiScribe™ T7 mRNA Kit with CleanCap® AG | New England Biolabs | — |
| Monarch® RNA Cleanup Kit | New England Biolabs | — |
| Lipofectamine™ MessengerMAX™ Transfection Reagent | Thermo Fisher Scientific | — |
| Opti-MEM™ I Reduced Serum Medium | Thermo Fisher Scientific | — |
| qScript™ cDNA SuperMix | Quantabio | — |
| PerfeCTa® SYBR® Green SuperMix | Quantabio | — |
| TRIzol™ Reagent | Invitrogen | — |
| CellMask™ Deep Red Plasma Membrane Stain | Thermo Fisher Scientific | — |
| CellMask™ Green Plasma Membrane Stain | Thermo Fisher Scientific | — |
| ProLong™ Gold Antifade Mountant with DAPI | Thermo Fisher Scientific | — |
| PVA DABCO Mounting Medium | Sigma-Aldrich | — |
| **Experimental models: organisms/strains** |  |  |
| Mouse: C57BL/6J | The Jackson Laboratory | Stock #000664 |
| Mouse: Arc<sup>-/-</sup> (AGN) | Wang et al., 2006 (gift; maintained at Cornell) | — |
| **Experimental models: cell lines** |  |  |
| DC2.4 murine dendritic-cell line | Millipore | SCC142 |
| B16-F10 melanoma | ATCC | CRL-6475 |
| EO771 breast cancer | Gift from Weiss Lab (Cornell) | — |
| **Oligonucleotides** |  |  |
| qPCR primers (*18S*, *Actb*, *Arc*, *SynA*, *Cd86*, *Il-2*, *Ifn-γ*, *Il-10*, *Tgfb*) | This study | Sequences in Supplementary Table 2 |
| **Recombinant DNA** |  |  |
| pGEX-6p1-GST-ArcFL | Addgene | #119877 |
| pcDNA3-EGFP | Addgene | #13031 |
| pcDNA3-Arc; pcDNA3-A5U-GFP | This study | Available on request |
| **Software and algorithms** |  |  |
| FlowJo (v10) | BD/FlowJo LLC | https://www.flowjo.com |
| GraphPad Prism (v9) | GraphPad | — |
| Gen5 (Cytation) | Agilent BioTek | — |
| ZEN blue | ZEISS | — |
| ImageJ / FIJI | NIH | https://imagej.nih.gov/ij |
| Living Image / Aura (IVIS analysis) | PerkinElmer / Spectral Instruments Imaging | — |
| Proteome Discoverer (v2.4), Sequest HT | Thermo Fisher Scientific | — |
| DIA-NN (v1.8) | Demichev et al. | — |
| G:Profiler (g:GOST) | University of Tartu | https://biit.cs.ut.ee/gprofiler |
| CFX Maestro (qPCR) | Bio-Rad | — |
| **Instruments** |  |  |
| Cytation 7 multimode imager | Agilent BioTek | — |
| Zeiss LSM 800 / 980 confocal microscopes | Carl Zeiss | — |
| IVIS Spectrum | PerkinElmer | — |
| NanoSight NS300 | Malvern Panalytical | — |
| NovoCyte Quanteon 4025 flow cytometer | Agilent | — |
| Bio-Rad CFX real-time PCR system | Bio-Rad | — |
| SkyScan 1276 micro-CT | Bruker | — |
| Ultrafiltration devices (100 kDa) | Amicon (MilliporeSigma) | — |
| TFF system, 300-kDa RC cassette | Millipore Pellicon Mini | — |

### Resource availability

#### Lead contact

Further information and requests for resources and reagents should be directed to and will be fulfilled by the Lead Contact, Shaoyi Jiang.

#### Materials availability

This study did not generate new stable cell lines. Plasmids encoding Arc and Arc 5′ UTR (A5U-GFP) are available from the Lead Contact upon request.

#### Data and code availability

All data supporting the findings of this study are available within the main text and Supplementary Information. Raw flow cytometry files, NTA data, and mass spectrometry datasets are available from the Lead Contact upon reasonable request. This study did not generate custom code.

### Experimental model and subject details

#### Mice

All animal procedures were performed at Cornell University’s Center for Animal Resources and Education (CARE) under approved IACUC protocols #2020-0037 and #2023-0101. Wild-type C57BL/6J mice (Jackson Laboratory) and Arc⁻/⁻ (AGN) mice ^27^ were maintained under specific pathogen-free conditions (12-h light/dark cycle, 22 ± 1 °C, 40-60% humidity) with ad libitum access to food and water. Mice of both sexes, 6-10 weeks old, were randomly assigned to treatment groups, with littermates balanced by sex where possible.

#### Cell lines

DC2.4 murine dendritic cells (Millipore, SCC142), B16-F10 melanoma cells (ATCC, CRL-6475), and EO771 breast cancer cells (gift from the Weiss Lab, Cornell) were maintained in DMEM or RPMI-1640 (ATCC) supplemented with 10% FBS (Gibco) at 37 °C in 5% CO₂. No antibiotics were used in stable cell culture. For in vivo experiments, donor cell strain was matched to the recipient mouse background.

#### Primary BM-DC/MΦ cultures

Primary bone marrow-derived DCs and macrophages (BM-DC/MΦs) were generated from C57BL/6J or Arc⁻/⁻ mice, as previously reported ^39^. Femurs and tibiae were disinfected in 70% ethanol, dissected, and flushed with HBSS or RPMI-1640 + 1% FBS using 25G (femur) or 27G (tibia) needles (20 mL per femur; 10 mL per tibia). Cells were filtered through a 70 µm strainer, centrifuged (180 ×g, 10 min), RBC-lysed in ice-cold eBioscience buffer (5 min), washed, and resuspended in complete RPMI-1640 containing 10% FBS, 2-mercaptoethanol, Pen-Strep, murine GM-CSF (20 ng/mL) and IL-4 (5 ng/mL). A total of 1 × 10⁷ cells were seeded per T25 flask in 5 mL. On day 2, half the medium was replaced with fresh medium containing 40 ng/mL GM-CSF and 10 ng/mL IL-4. On day 3, the medium was fully replaced with 20 ng/mL GM-CSF and 5 ng/mL IL-4. On day 6, all adherent, loosely attached, and suspension cells were harvested with warm DPBS and pooled as BM-DC/MΦ donors.

### Method details

#### EV production and donor cell transfection

Donor DC2.4 cells or BM-DC/MΦs were transfected with capped, polyadenylated Arc and A5U-GFP mRNAs using Lipofectamine MessengerMAX in Opti-MEM according to the manufacturer’s protocol. A5U-GFP served both as a stabilizing RNA element and a fluorescent reporter for transfection efficiency. Live-cell epifluorescence imaging (Cytation 7) was used to monitor GFP expression and optimize the time window for peak VLV production. EVs were produced in serum-free Opti-MEM to increase vesicle yield and reduce serum-derived contamination. EV preparations were freshly generated for each in vivo or in vitro experiment; pilot studies indicated that long-term storage markedly decreased vesicle integrity and function.

#### EV fluorescent labeling and NTA

Freshly isolated EVs were quantified by NTA and adjusted to 1 × 10¹¹ particles/mL. For in vivo tracking, EVs were stained with CellMask™ Deep Red (1:5000 in 0.1 µm-filtered DPBS; 10 min, RT, gentle rocking, protected from light). Unbound dye was removed by 5-6 ultrafiltration washes at 4 °C (100-kDa cutoff) with filtered DPBS until a mock DPBS control processed in parallel appeared completely blank under identical fluorescence settings. Preparations in which the DPBS blank remained fluorescent were discarded and re-stained with fresh dye at lower dye concentration or higher EV density to avoid dye micelles. Labeled EVs were finally resuspended at 1 × 10¹² particles/mL in low-bind tubes and stored at 4 °C until injection.

For fluorescent NTA, labeled EVs were analyzed on a NanoSight NS300 using 488- or 532-nm lasers and appropriate emission filters. Light-scatter mode was used to obtain total particle counts, while fluorescence mode distinguished labeled EVs from background. Negative controls guided camera settings to exclude non-fluorescent noise. Particle size distributions and concentrations were calculated from Brownian motion trajectories.

#### Mouse vaccination and tumor challenge

Engineered vesicle vaccines (B16⁺ DEVs or VLVs) were administered intravenously via the retro-orbital sinus once weekly for four consecutive weeks. Injections alternated between eyes to minimize local irritation. Each dose contained 50 µL of EV suspension with up to 6 × 10¹⁰ particles per mouse. EV numbers were determined by NTA, and batch-specific particle counts and dosing records are summarized in Supplementary Table 1. Control animals received volume-matched sterile DPBS. Mice were monitored until full recovery from anesthesia and then daily for signs of distress, weight loss, or injection-site inflammation; body weight and activity/grooming scores were recorded weekly. No treatment-related mortality or severe toxicity was observed.

Tumor measurements began 5 days after B16-F10 injection and continued daily. Tumor dimensions were measured with digital calipers in a double-blind fashion by a team of five researchers; the treatment group identity was concealed during measurement. For noninvasive 3D tumor assessment, micro-CT scans (SkyScan 1276) were acquired under anesthesia. Tumor volumes were segmented in dedicated software by delineating tumor boundaries across slices and summing slice volumes to derive growth curves.

### Microscopy and *in vivo* imaging

#### Epifluorescence live-cell imaging

Cells were plated on glass-bottom dishes and imaged on a Cytation 7 imager with temperature and CO₂ control. GFP/A5U-GFP expression and morphology were monitored at multiple time points post-transfection. DAPI staining was used when needed to quantify nuclei and cell density. Gen5 software was used for acquisition and analysis, and these readouts informed optimal EV collection timing.

#### Confocal microscopy

High-resolution imaging of cells and tissues was performed on a Zeiss LSM 980 with 405, 488, 561, and 640-nm lasers and 5×, 10×, 20×, and 60× objectives. Optical sections were acquired at 1 Airy unit pinhole with 2-4× frame averaging. Images were processed in ZEN blue and analyzed in ImageJ/FIJI for intensity, co-localization, and EV distribution.

#### Immunocytochemistry and immunohistochemistry

For ICC, cells were fixed in 4% PFA (15 min), permeabilized with 0.1% Triton X-100, and blocked in 3% normal goat serum prior to overnight incubation with primary antibodies (4 °C). Fluorophore-conjugated secondary antibodies were applied and samples mounted in ProLong™ Gold (with DAPI) or PVA DABCO. For IHC, cryosections (10-30 µm) or paraffin sections were rehydrated, subjected to antigen retrieval where required, and stained with primary antibodies against EV or cellular markers followed by fluorescent secondaries. Imaging parameters were kept constant within experiments to allow quantitative comparison of staining intensity.

#### IVIS imaging

For organ-level EV distribution, mice received CMDR-labeled EVs and were perfused trans-cardially with PBS under deep anesthesia 6 h post-injection to reduce blood background. Organs were imaged on an IVIS Spectrum using CMDR-appropriate filter sets. Unlabeled controls were included each session to correct for autofluorescence. Radiance (photons s⁻¹ cm⁻² sr⁻¹) within regions of interest was quantified with Living Image or Aura software and exported to ImageJ for normalization and statistical analysis.

### Immune-response analysis

#### Cell collection

At designated time points, mice were euthanized, and spleens and lymph nodes were harvested aseptically. Splenocytes were prepared by mechanical disruption through a 70 µm strainer, RBC-lysed, washed, and resuspended in Opti-MEM or flow buffer. Lymphocytes from pooled lymph nodes were obtained by mechanical disruption and sequential filtration through 70 µm and 40 µm strainers.

#### Cell flow cytometry

Single-cell suspensions were stained with a panel including Live/Dead dye, CD45, CD3, CD4, CD8, CD44, CD62L, and intracellular IFN-γ, along with additional markers as required. Staining was performed in standard flow buffer with Fc blocking. Samples were acquired on a NovoCyte Quanteon 4025 and analyzed in FlowJo v10. T cell subsets (naïve, central memory, effector/effector memory) were defined based on CD44/CD62L expression.

#### EV flow cytometry

Purified EVs from DC2.4 cells or primary BM-DC/MΦ donors were analyzed at single-vesicle resolution using a NovoCyte Quanteon 4025 flow cytometer (Agilent), operated at the lowest stable flow rate (5 μL/min) to minimize swarm events. All buffers were 0.1 μm-filtered. EVs were stained in DPBS/F68 using the same antibody panel used for donor-cell phenotyping, CD45, CD11c, CD40, CD80, CD86, MHC-I (H-2Kᵇ), MHC-II (I-A/I-E), and H-2Kᵇ:SIINFEKL, along with Live/Dead and compensation controls.

To remove unbound antibodies, stained EVs were cleaned using the EXODUS H-600 automatic exosome isolation system (Exodus Bio), which employs dual-membrane nanofiltration with negative-pressure oscillation and ultrasonic harmonic pulses. This step efficiently eliminates free fluorophores and antibody aggregates while retaining intact vesicles, enabling reliable detection of fluorescently labeled EVs.

Controls on every run included (1) dye-only EVs, (2) no-stain EVs, (3) FMO controls, (4) detergent-lysed EVs + stain to identify nonspecific binding, and (5) serial dilutions to verify linear detection ranges. To ensure data quality, unstable time segments and multiplet events were removed through initial gating.

Vesicles were identified as ExoBrite⁺ (true EV membrane stains) CD45⁺ CD11c⁺ singlets. MFI, percent-positive fractions, and contour-density distributions were analyzed in the NovoCyte software. Surface-marker density was compared between EVs and their donor DCs using MFI and surface-normalized metrics (Enrichment Index, EI), and population composition was quantified by gating high-expression subsets (CD40^hi^, CD86^hi^, MHC-I^hi^, MHC-II^hi^, H-2Kᵇ:SIINFEKL⁺). All EV flow cytometry assays were performed with matched reagent lots and identical thresholding parameters within each experiment.

#### ELISA

At weeks 3, 4, 6, 9, and 13 after the first immunization, blood was collected by cheek bleed. Serum IgG, IgM, and IgA against Ova were quantified by ELISA. Plates were coated with Ova antigen, blocked (10% normal goat serum in SuperBlock), incubated with serially diluted serum, followed by HRP-conjugated secondaries and TMB substrate. Reactions were stopped with sulfuric acid and read on a Cytation 7; titers were calculated from standard curves or endpoint dilution.

#### RT-qPCR

Total RNA was extracted with TRIzol/BCP, DNase-treated, and purified (PureLink). cDNA was synthesized from 2 µg RNA using the High-Capacity cDNA Reverse Transcription Kit. qPCR was performed with PowerUp SYBR Green Master Mix using primers for cytokines and virus-like components (primer sequences in Supplementary Table 2). Cycling conditions were 95 °C for 2 min, followed by 40 cycles of 95 °C for 15 s and 60 °C for 1 min. Melt-curve analysis confirmed primer specificity. Relative expression was calculated using ΔCt or ΔΔCt with 18S and Actb as reference genes.

#### ELISpot

To assess antigen-specific IFN-γ-secreting cells, ELISpot plates were coated with anti-IFN-γ capture antibodies. Splenocytes from boosted mice were restimulated with Ova MHC-I or MHC-II peptides or B16-F10 EVs. After incubation, plates were washed, incubated with detection antibodies and substrate, and developed until spots were visible. Spots were counted on a Cytation 7 and reported as spot-forming units per 10⁶ cells.

#### *Ex vivo* cytotoxicity and splenocyte-tumor co-culture

BM-DC/MΦs from WT and Arc⁻/⁻ donors were prepared as above. A subset of WT BM-DC/MΦs was transfected with Arc and A5U mRNAs to generate Arc-overexpressing donors; WT and Arc⁻/⁻ cells without transfection served as controls. All donor groups were exposed to B16-F10 EVs for 24 h to provide melanoma antigens, and resulting vesicles were purified and quantified by NTA.

C57BL/6J mice were vaccinated intravenously with vesicles from: (1) Arc⁺ A5U⁺ BM-DC/MΦs, (2) WT BM-DC/MΦs, or (3) Arc⁻/⁻ BM-DC/MΦs. Eight days later, spleens were harvested and splenocytes prepared by mechanical disruption and RBC lysis. Splenocytes were labeled with CFSE (5 µM, 10 min, 37 °C) and washed. B16-F10 targets were pre-stained with NucSpot 650. Effector and target cells were co-cultured in 96-well plates at ∼200:1 E:T (0.6 × 10⁶ splenocytes + 3 × 10³ targets per well in 100 µL Opti-MEM) with 0.5 µg/mL PHA-P to maintain viability without strong nonspecific activation. Plates were incubated 24-48 h at 37 °C, 5% CO₂. Controls included targets-only, effectors-only, PBS-splenocytes (± PHA-P), and medium-only wells.

After 24 h, non-adherent cells were gently removed and wells washed with warm Opti-MEM. Adherent tumor cells were imaged on a Cytation 7 using GFP (CFSE) and Cy5 (NucSpot) channels. Effector-target interaction frequency was quantified as the percentage of NucSpot⁺ tumor cells in contact with CFSE⁺ splenocytes across five random high-power fields per well. All conditions were run in triplicate from independent mice.

### Mass spectrometry sample preparation and LC-MS/MS data acquisition

#### Protein extraction and SDS-PAGE

EVs from Arc-overexpressing or control DCs were purified by TFF and ultrafiltration. Vesicles were lysed in IP buffer with detergents and protease inhibitors, subjected to one freeze-thaw and vortexing on ice, then cleared by centrifugation. Protein concentration was determined (Pierce 660 assay). Equal protein amounts were denatured in SDS with β-mercaptoethanol, separated by SDS-PAGE, and stained (SimplyBlue or silver). Bands of interest were excised for in-gel digestion.

#### In-gel digestion

Gel pieces (∼2 mm) were washed (water; 50 mM ammonium bicarbonate/50% ACN; 100% ACN), dried (SpeedVac), reduced with 10 mM DTT (100 mM ammonium bicarbonate, 1 h, 60 °C), alkylated with 55 mM iodoacetamide (dark, 45 min, RT), washed again, dried, and rehydrated with trypsin (10 ng/µL, 50 mM ammonium bicarbonate/10% ACN) on ice. Additional buffer was added and digestion proceeded at 37 °C for 16 h. Peptides were extracted sequentially with 2% FA, 50% ACN/5% FA, and 90% ACN/5% FA, pooled, dried, and reconstituted in 2% ACN/0.5% FA before filtration (0.22 µm) and nanoLC-MS/MS.

#### Orbitrap Fusion analysis

Peptides were analyzed on an Orbitrap Fusion Tribrid coupled to a Dionex UltiMate 3000 RSLCnano. Samples were trapped on a PepMap C18 column and separated on a C18 analytical column with a 90-min gradient (5-35% ACN/0.1% FA, followed by 90% ACN wash). The Orbitrap was run in positive mode with 1.2 kV spray voltage and 275 °C source. Data-dependent acquisition used MS1 scans at 120,000 resolution (m/z 300-1600) followed by 3 s “Top Speed” CID MS/MS in the ion trap (normalized collision energy 30%). Dynamic exclusion was 50 s (±10 ppm).

#### timsTOF Pro / diaPASEF analysis

For DIA, peptides (100 ng) were separated on a NanoElute LC coupled to a timsTOF Pro via CaptiveSpray, using a 21-min gradient (2-30% ACN/0.1% FA; then 30-95% ACN). Data were acquired in diaPASEF mode with 16 m/z and ion mobility windows, 1.5 kV spray, 180 °C transfer tube, and m/z 100-1700. Collision energy was ramped from 20 to 59 eV across the ion mobility range.

#### Protein identification and quantification

Orbitrap DDA files were searched in Proteome Discoverer 2.4 with Sequest HT against the Mus musculus NCBI RefSeq database (∼28,230 entries) plus an EV database including Ova sequences. Precursor tolerance was 10 ppm, fragment tolerance 0.6 Da. Static modification: carbamidomethyl C; variable modifications: M oxidation, N/Q deamidation, protein N-terminal acetylation, M-loss, and M-loss + acetylation. Percolator FDR was set to 1% at the peptide level. Label-free quantification used the Minora Feature Detector; peptide abundances were aligned and summed as unique + razor peptides per protein. diaPASEF data were processed in DIA-NN v1.8 using an in-house spectral library built from the UniProt-SwissProt Homo sapiens database (Taxon ID 9606, downloaded 01/20/2023, 20,404 entries). Cysteine carbamidomethylation was set as a fixed modification; methionine oxidation and N-terminal acetylation as variable. FDR was controlled at <1% at peptide and protein levels.

#### Quantification and statistical analysis

Unless otherwise stated, data are presented as mean ± SEM from biological replicates (individual mice or independent cultures). Group comparisons were performed in GraphPad Prism using unpaired two-tailed t-tests, one-way ANOVA or two-way mixed-effects ANOVA with appropriate post hoc multiple-comparison tests (typically Dunnett’s), as appropriate for the design. Survival curves were analyzed by log-rank (Mantel-Cox) tests. P < 0.05 was considered statistically significant.

#### Disclosure of AI assistance

ChatGPT was used after drafting for grammar and style refinement only. It did not create original content, perform analyses, or generate data. The authors take full responsibility for the accuracy and integrity of the work.

## Acknowledgements

S.J. acknowledges start-up support from Cornell University, including Robert Langer ’70 Family and Friends Professorship, Cornell NEXT Nano Initiative and Cornell Engineering’s inaugural Sprout Award. We thank the Proteomics and Metabolomics Facility of Cornell University for providing the mass spectrometry data and NIH SIG grant 1S10 OD017992-01 support for the Orbitrap Fusion mass spectrometer. We thank Andy Yu-Wei Chang and the Weiss lab at Cornell for their kind gift of EO771 cells. We thank Prof. Kuan Hong Wang, University of Rochester Medical Center, for his kind gift of Arc^‒/‒^ AGN mice. We thank Tina Abratte, Teresa Porri and the BRC Imaging Facility at the Cornell Institute of Biotechnology (RRID:SCR_021741) for imaging experiments, with NIH S10OD025049 for the IVIS-Spectrum optical imager and the SkyScan 1276 mouse CT in Cornell’s BRC Imaging Facility. We thank Emily Silvela, Elizabeth S. Moore, Malik J. Scales, Trevor Totman, Erica Feldman, Faith Burgus, at Cornell CARE for their service and advice on caring for animals. We thank Cornell IACUC for their help composing and managing animal protocols.

## Author contributions

W. Gu and S. J. conceptualized the work. W. Gu, R. L., H. G., D. A., S. C., I. T., A. L., A. S., V. U., N. E., S. L., G. Q., Y. Z., Y. Y., E. G., J. L., and E. S. acquired the data. W. Gu, R. L., H. G., and D. A. contributed to data analyses and to the interpretation of the results. W. Gu, R. L., H. G., C. C., Q. Y. and S.J. provided advice, support and supervision. W. Gu wrote the manuscript. S.J. edited the manuscript.

## Declaration of Interests

S. J. and W. Gu are authors of two provisional patent applications related to this work filed by Cornell University. All other authors declare no competing interests.

