## Supplemental information for "Repurposed endogenous virus-like vesicles mediate dendritic cell long-range antigen presentation and T cell activation for enhanced cancer vaccination"

**Supplementary figures**

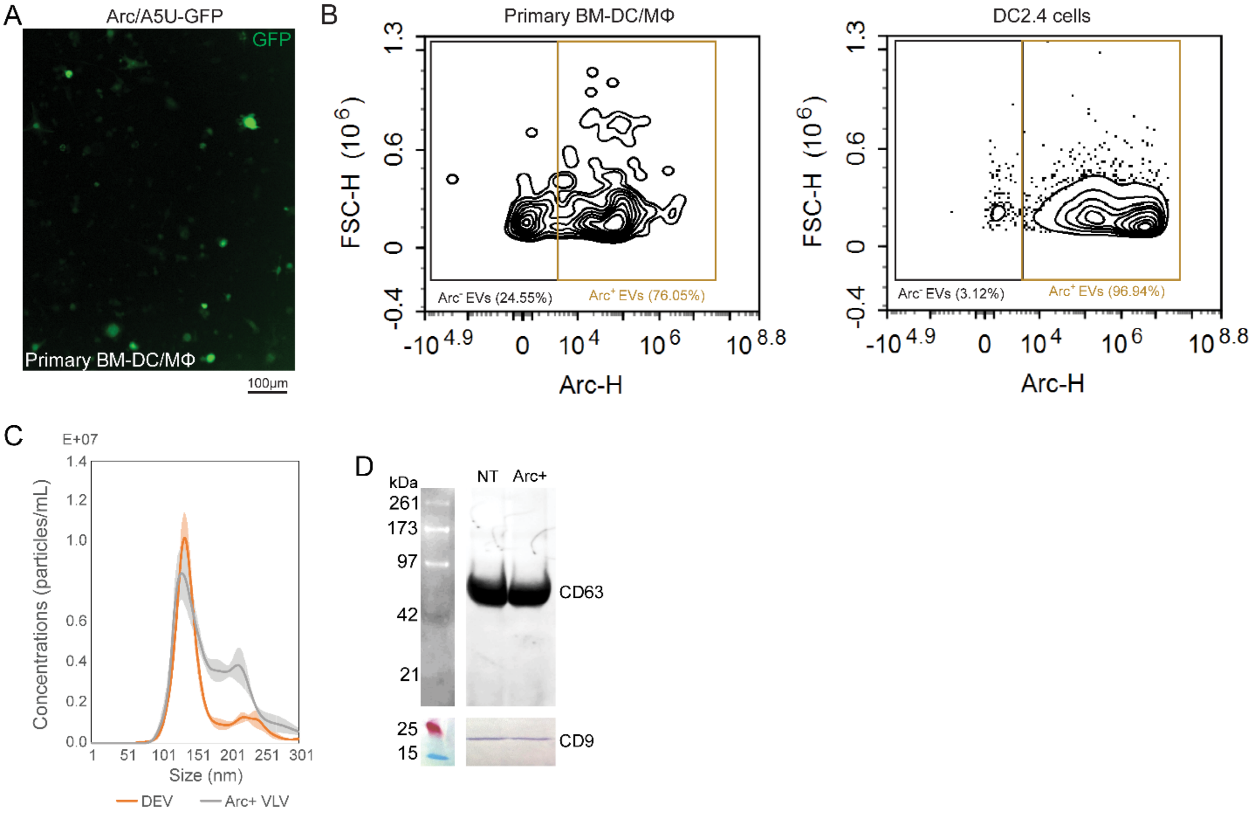


**Figure S1.** *Characterization of engineered EVs.* Each EV batch undergoes standardized characterization, including assessment of donor-cell reporter expression and NTA, with western blot and single-vesicle flow cytometry performed as needed. **(A)** Donor-cell RNA-liposome transfection efficiency: Primary BM-DC/MΦs were transfected with Arc and A5U-GFP RNAs, assessing transfection efficiency. **(B)** Single-vesicle flow cytometry: ~70-80% of EVs are Arc$^+$ when engineered from primary BM-DCs/MΦs, and up to ~97% are Arc$^+$ when produced from the stable DC2.4 line. Notably, very high Arc expression does not necessarily confer improved functionality; excess Arc tends to be shed in exosomal fractions that lack functionality. **(C)** NTA: This panel shows representative size-distribution profiles comparing unmodified DEVs and engineered Arc$^+$ VLVs. Engineered Arc$^+$ VLVs exhibit a distinct, larger peak relative to the major peak of unmodified DEVs. **(D)** Western blot for EV markers: Immunoblotting confirms the presence of canonical EV proteins in all samples.

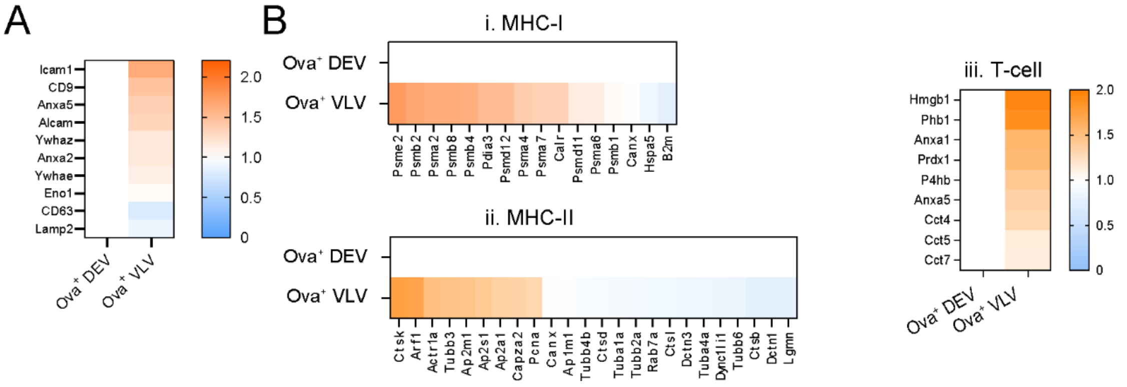


**Figure S2.** *Proteomic enrichment of immune-regulatory and antigen-presentation proteins in Ova⁺ VLVs compared with Ova⁺ DEVs.* **(A)** Heat map of label-free MS quantification showing increased abundance of adhesion and budding proteins (ICAM1, CD9, Annexin A5, ALCAM) and decreased endosomal markers (CD63, LAMP2) in Ova⁺ VLVs relative to Ova⁺ DEVs. **(B)** Functional categorization of upregulated proteins in Ova⁺ VLVs versus Ova⁺ DEVs. (i) MHC-I-associated proteins (e.g., Psme2, Psmb8, Tap1, H2-K1, B2m) were enriched, suggesting enhanced class I antigen presentation. (ii) MHC-II-associated proteins (e.g., Ctss, Cd74, H2-Aa, H2-Ab1) were maintained or modestly increased, indicating preserved class II presentation. (iii) T-cell–interacting proteins (e.g., Hmgb1, Phb1, Annexin A1/A5, Cct4/5/7) were upregulated, supporting enhanced immunogenic potential of VLVs. Data represent average relative intensities of two biological replicates × two technical replicates ($\log_2$ fold change, VLV/DEV).

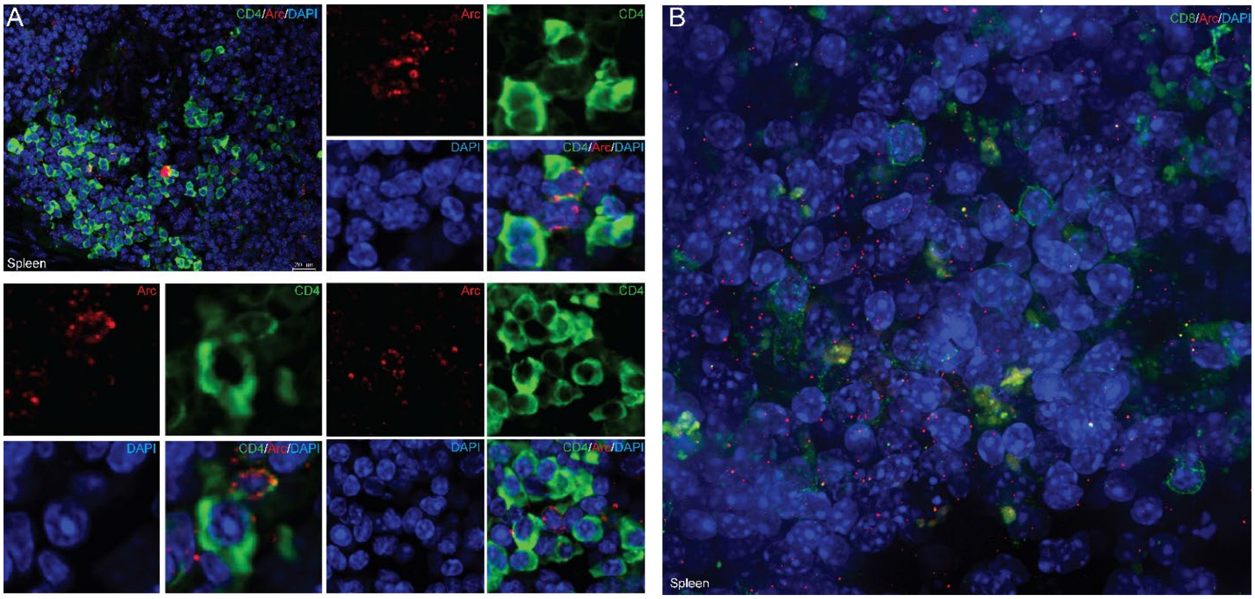


**Figure S3.** *Analysis of Arc⁺ vesicle distribution in spleen tissue***.** This figure provides an expanded view of spleen tissue, offering a broader perspective than the main figure by illustrating the widespread presence of both $Arc^+$ cells and $Arc^+$ EVs within the spleen. It includes detailed zoom-in regions that emphasize variations in CD4 and CD8 expression levels among different cell populations, allowing for a finer examination of the interaction dynamics between $Arc^+$ EVs with varying degrees of CD4 and CD8 expression.

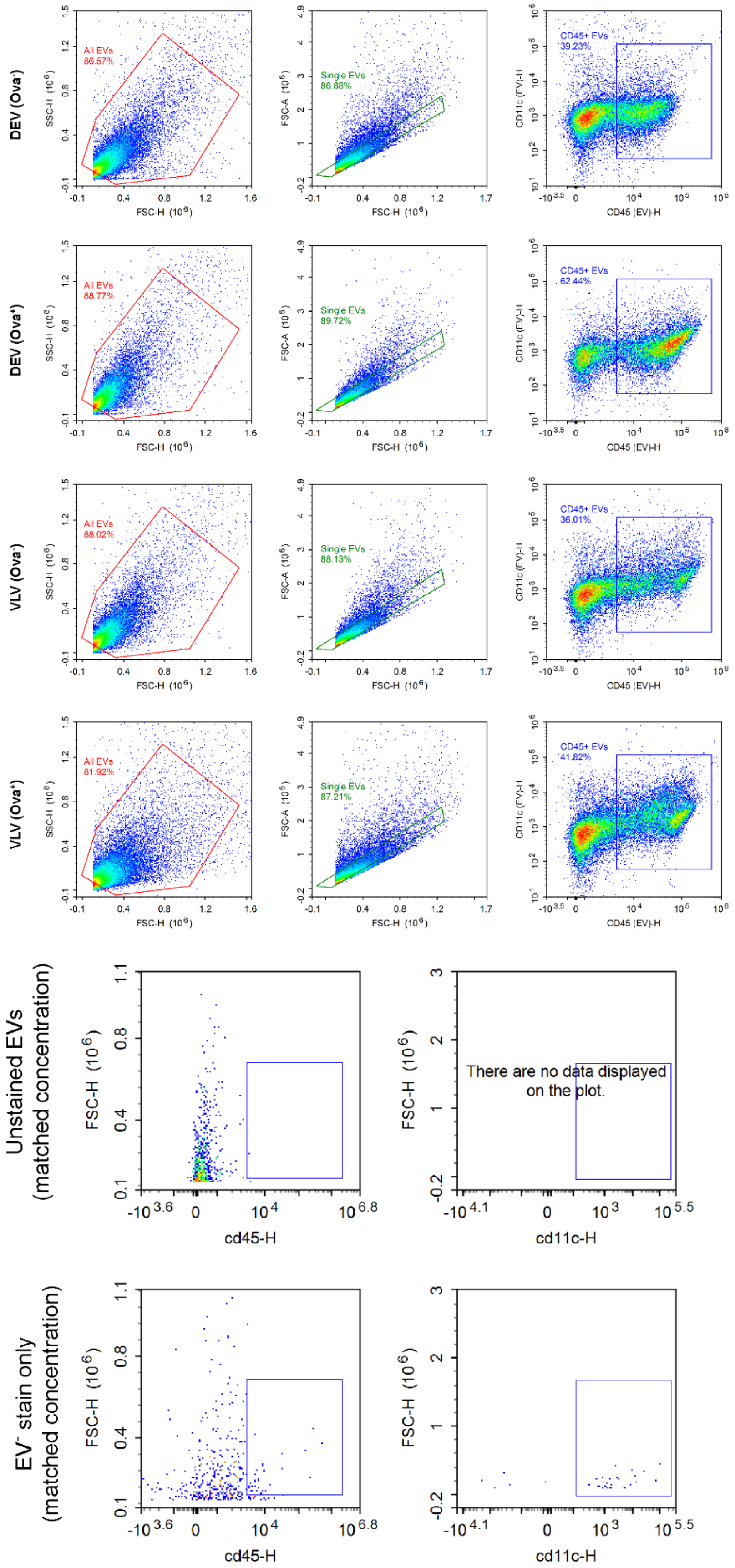


**Figure S4.** *Gating strategy for DEVs.* Representative flow cytometry plots showing sequential gating of DEVs and VLVs. Events were first gated by forward scatter (FSC-H) versus side scatter (SSC-H) to define the EV population (red gate, left column), followed by selection of singlet EVs using FSC-A versus FSC-H (green gate, middle column). Within the singlet population, $CD45^+/CD11c^+$ EVs were identified (blue gate, right column) to define DEVs. Percentage of total and gated EVs are indicated on each plot. Negative controls (unstained EVs at matched concentrations and dye-only without EVs) showed no significant $CD9^+/CD45^+/CD11c^+$ populations.

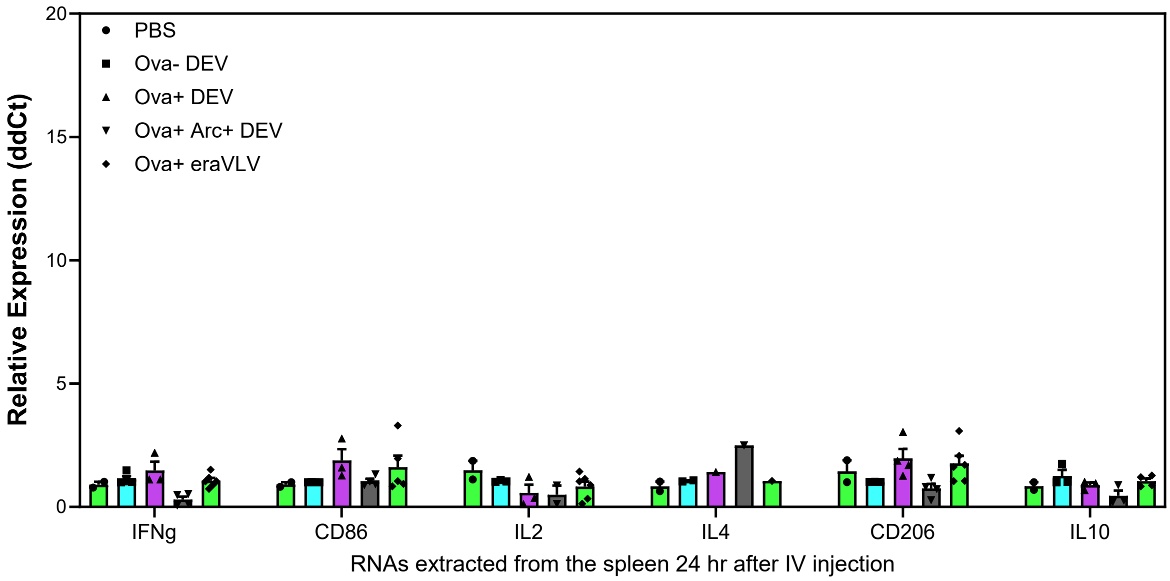


**Figure S5.** *RT-qPCR analysis of immune-related gene expression in the spleen 24 hours after DEV/VLV treatment***.** Expression was normalized to Gapdh and to the control group using the $2^{\Delta\Delta Ct}$ method. Data are presented as mean ± SEM, $N = 2$-6 biological replicates. Individual data points are shown, and no significant differences were observed among groups.

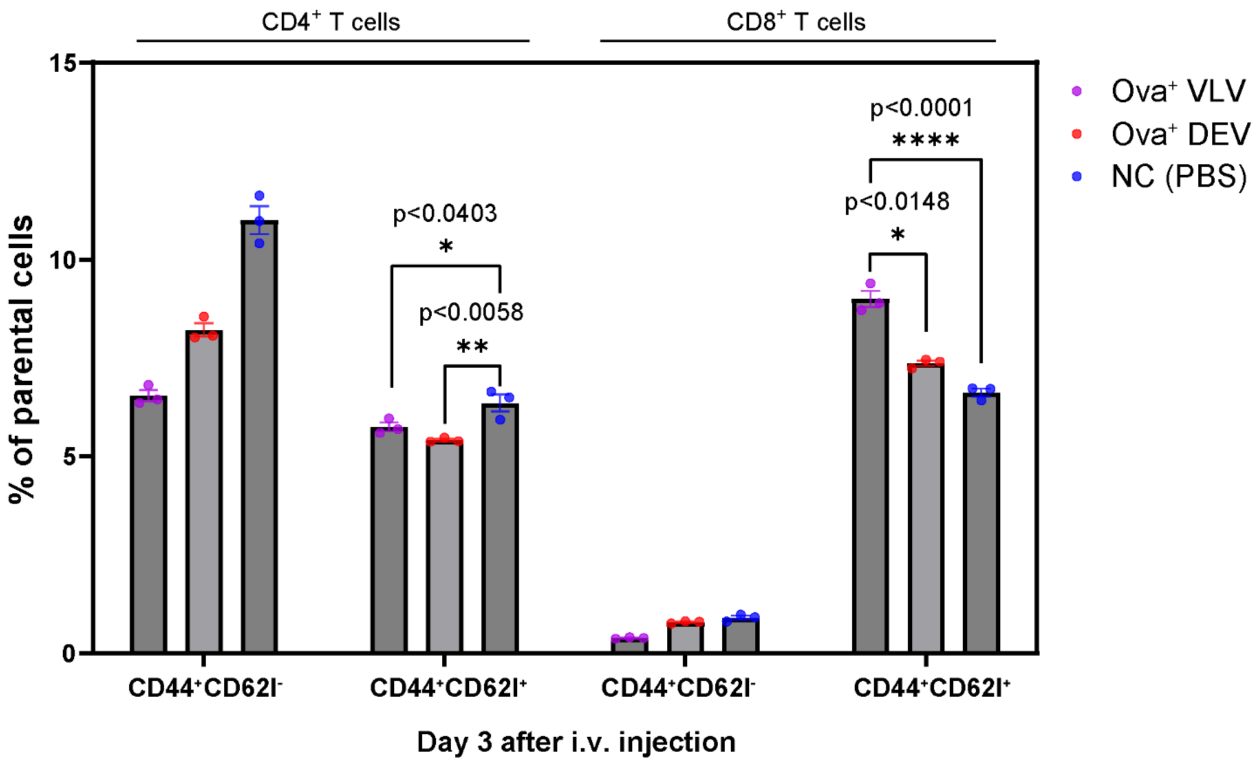


**Figure S6.** *Flow cytometry analysis of T cell populations in splenocytes isolated from C57BL/6 mice treated with WT or Arc KO DEVs presenting the model Ova antigen.* 3 days post-administration, Arc$^{+}$ VLV derived from DC2.4 donor cells significantly increased CD8$^{+}$ CD44$^{hi}$ CD62L$^{hi}$ T cells compared with both PBS and WT DEV controls, whereas CD4$^{+}$ CD44$^{hi}$ CD62L$^{hi}$ frequency decreased in both Arc$^{+}$ VLV and WT DEV groups. 3 biological replicates.

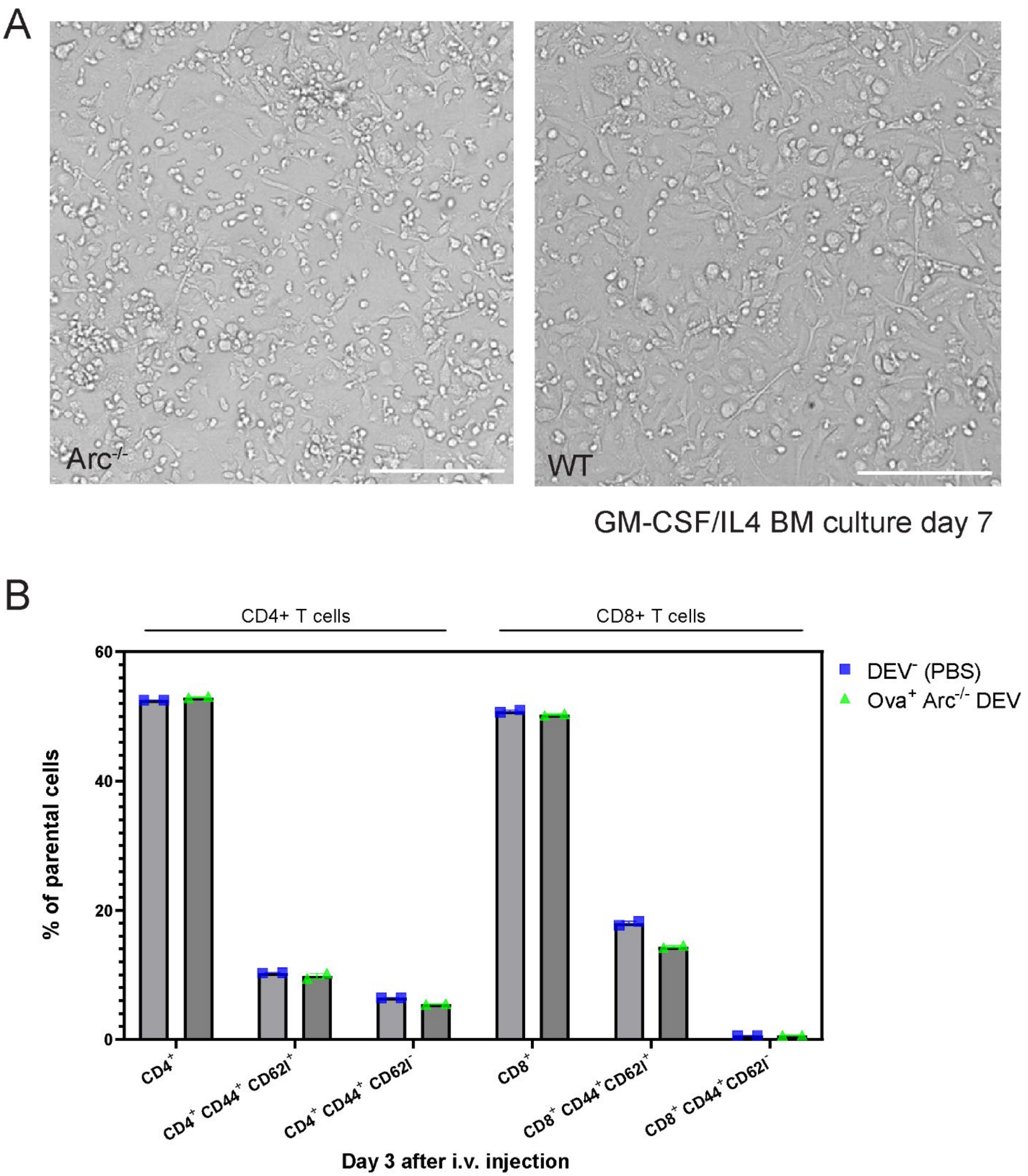


**Figure S7.** *Arc is necessary for APC morphology and vesicle-mediated T-cell activation.* **(A)** Morphology of primary BM-derived DC/MΦ from WT and $Arc^{-/-}$ mice on day 6 of differentiation. $Arc^{-/-}$ cells appeared smaller and less uniform than WT. Representative bright-field images; scale bars indicated. **(B)** Flow cytometry analysis of T-cell populations in splenocytes isolated from C57BL/6 mice treated with $Arc^{-/-}$ DEVs presenting the model OVA antigen, 3 days post-administration. Events were gated singlets → live → $CD45^{+}$ → $CD3^{+}$ → $CD4^{+}$, $CD8^{+}$, CD44 and CD62L. $OVA^{+}$ $Arc^{-/-}$ DEVs did not induce significant changes in $CD3^{+}$, $CD4^{+}$, $CD8^{+}$, or $CD44^{hi}$ $CD62L^{hi}$ frequencies compared to PBS. Mean ± SEM with individual data points (n=2).

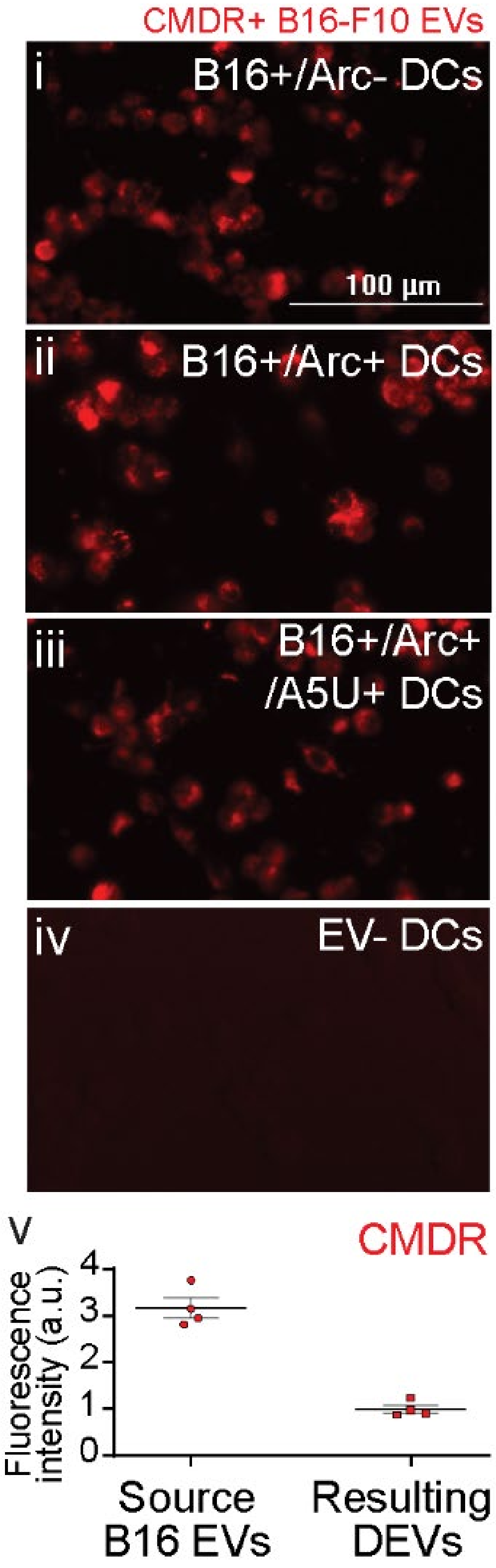


**Figure S8.** *Uptake of B16-F10 EVs by donor DCs and confirmation of minimal carry-over into DC-EV preparations.* B16-F10 EVs were labeled with CMDR, a lipid membrane dye. (i–iv) Fluorescence imaging of donor DC cultures shows efficient uptake of CMDR-labeled B16-F10 EVs, providing visual evidence of melanoma antigen incorporation. (iv) As a negative control, CMDR was added to EV-free DPBS and processed through the same ultrafiltration steps as B16-F10 EVs before being applied to DCs, confirming the specificity of the labeling method. (v) Purity of the collected DEV vaccine samples was assessed by CMDR fluorescence intensity, showing that DEVs used in subsequent experiments have minimized B16-F10 EV carry-over. Data represent mean ± SEM from four biological replicates. The residual fluorescence observed in DEVs reflects CMDR dyes incorporated into DC membranes following B16-F10 EV uptake and subsequently released on DC-derived EVs.

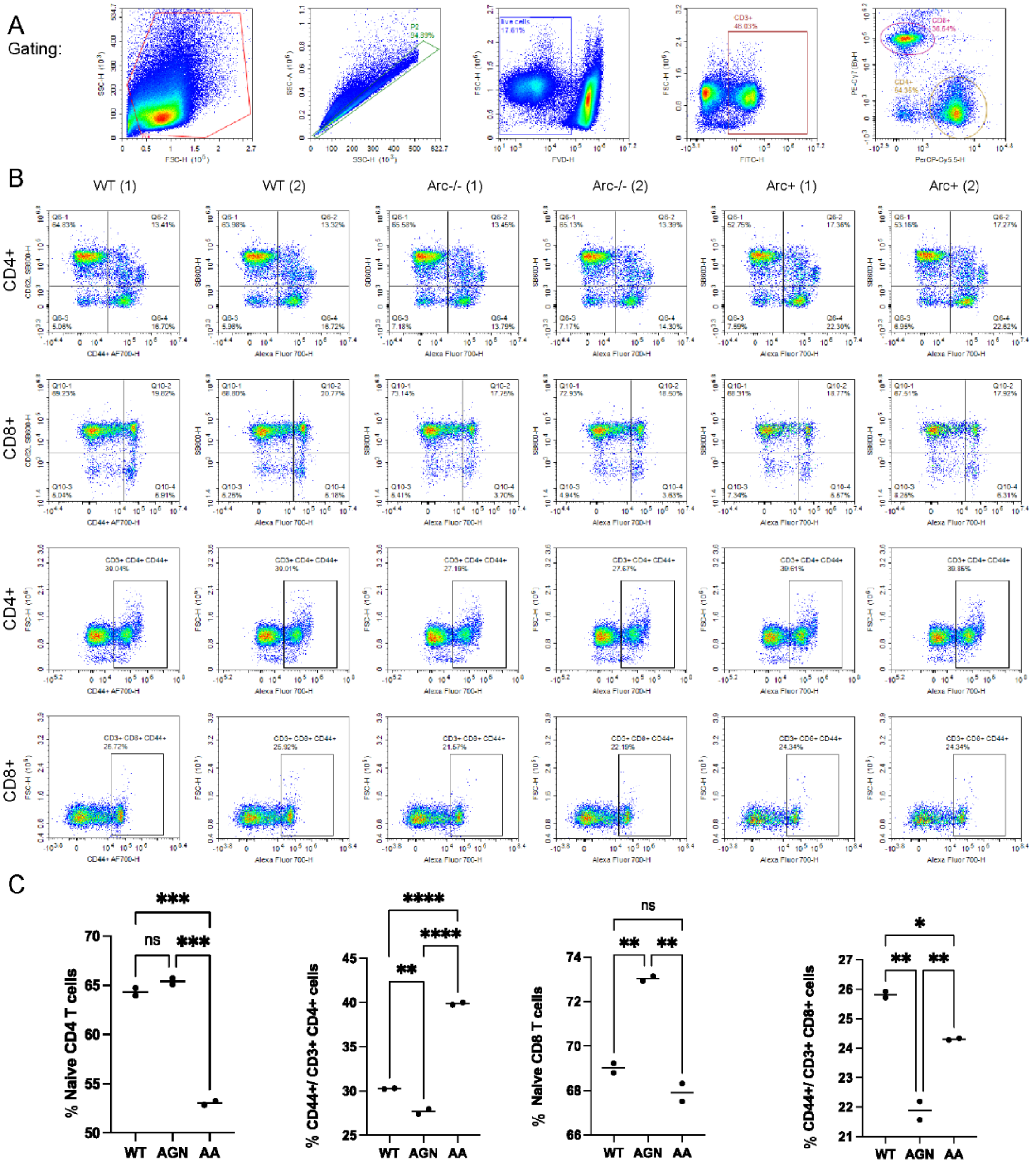


**Figure S9.** *Flow cytometry gating and memory-phenotype enrichment after restimulation (day-8 post vaccination).* **(A)** Representative flow cytometry workflow for splenocytes harvested 8 days post-vaccination and restimulated *ex vivo* with B16-F10-derived EVs plus co-purified proteins (6 h in serum-free OptiMEM followed by 6 h with a protein-transport inhibitor). Events were gated as singlets → live → $CD45^+$ → $CD3^+$→ $CD4^+$ or $CD8^+$, then profiled for memory/activation markers (CD44, CD62L). Fluorescence-minus-one (FMO) defined thresholds. **(B)** Frequencies of T cells populations across treatment groups. $Arc^+$ VLV vaccination increased $CD4^+CD44^+CD62L^-$ and $CD8^+CD44^+CD62L^-$ frequencies relative to both WT and $Arc^{-/-}$ groups. $Arc^+$ VLV vaccination increased $CD4^+CD44^+$ but not $CD8^+CD44^+$ frequencies relative to the WT control, whereas the $Arc^{-/-}$ group led to derecreased frequcnies of both $CD4^+CD44^+$ and $CD8^+CD44^+$ populations. (C) $Arc^+$ VLV treatment resultsed in significantly increased $CD4^+CD44^+$ frequencies relative to WT DEV controls, whereas $Arc^{-/-}$ DEVs led to significantly lower $CD4^+CD44^+$ and $CD8^+CD44^+$frequencies compared with both WT and $Arc^+$ groups. Data are shown as mean ± SEM with individual data points (each point = one mouse). Statistics: one-way ANOVA with Tukey's multiple-comparisons test; significance is annotated on the figure (*$p < 0.05$, **$p < 0.01$, ***p < 0.001, ****$p < 0.0001$).

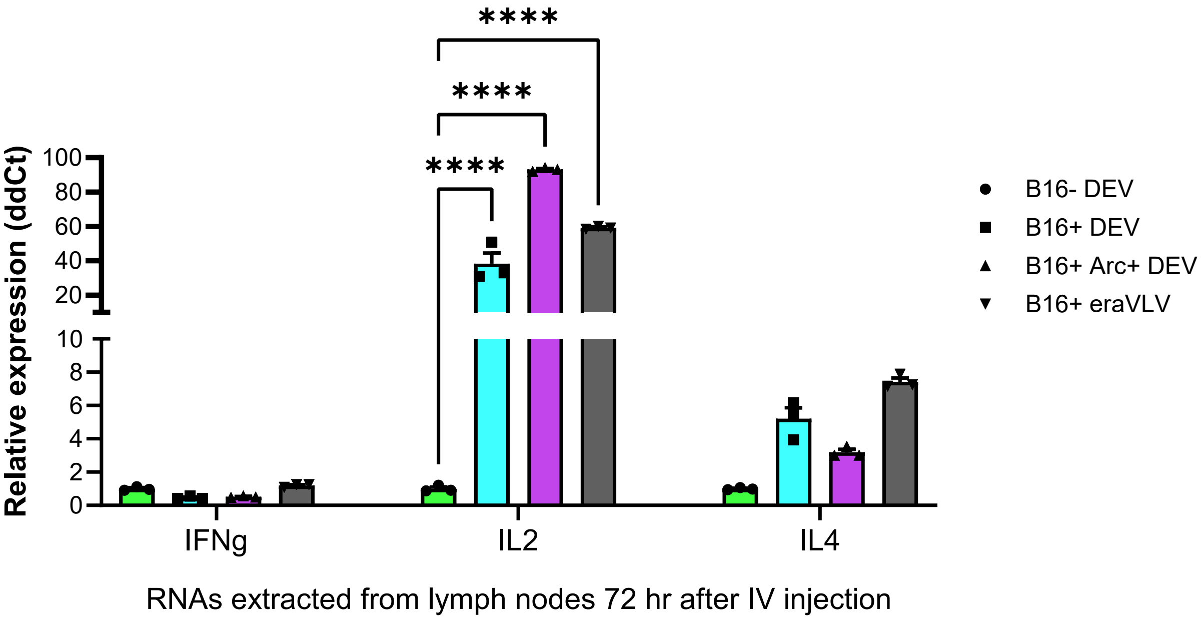


**Figure S10.** *RT-qPCR analysis of immune-related gene expression in pooled lymph nodes* (6 cervical and 4 axillary/brachial per mouse) 72 h after DEV/VLV treatment. Expression was normalized to Gapdh and to the control group using the $2^{\Delta\Delta Ct}$ method. Data are mean ± SEM (n = 3 biologically independent mice); individual data points shown. Statistics: one-way ANOVA; ****$P < 0.0001$.

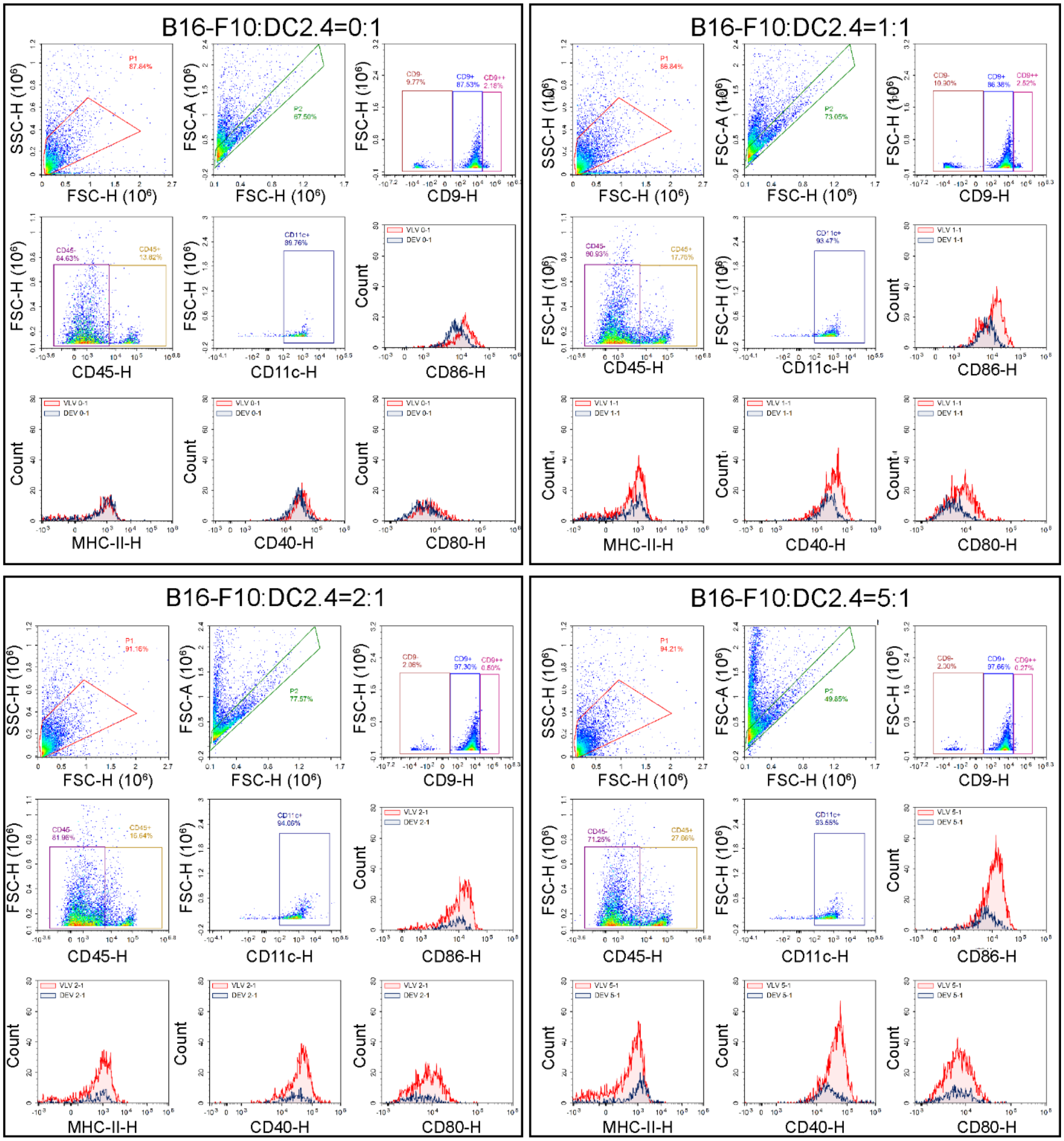


**Figure S11.** *EV flow cytometry characterization of B16-F10-primed DC2.4-derived DEVs and VLVs at different tumor-to-DC2.4 ratios.* Representative flow cytometry plots showing gating strategy and marker expression for DEVs and VLVs generated using B16-F10:DC2.4 input ratios of 0:1, 1:1, 2:1, and 5:1. Events were first gated on singlet $CD9^{+}$ $CD45^{+}$ and $CD11c^{+}$ EVs, followed by analysis of CD86, CD80, CD40, and MHC-II expression. As the relative number of B16-F10 cells increased, both co-stimulatory molecule counts and peak intensities (CD86, CD80, CD40) rose, while $MHC\text{-}II^{+}$ particle counts also increased with higher B16-F10 input.

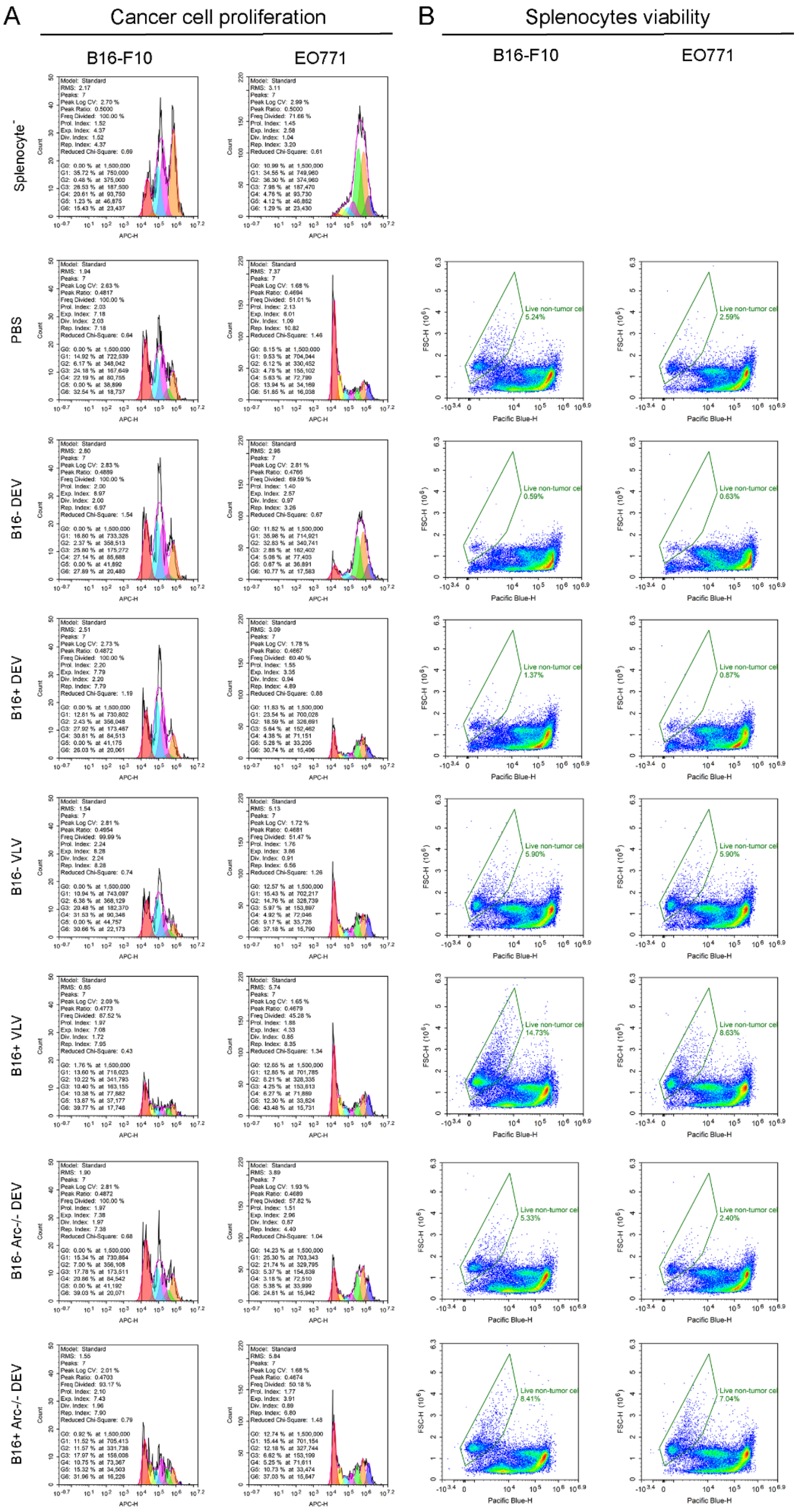


**Figure S12.** *Ex vivo tumor cell proliferation and responder T cell viability following DEV/VLV vaccination.* **(A)** Cancer cell proliferation. CellTracker-labeled B16-F10 (left) or EO771 (right) tumor cells were co-cultured for 48 h with splenocytes harvested from mice vaccinated with the indicated formulations. Representative generation histograms (APC-H channel) show progressive dye dilution corresponding to tumor cell divisions. The B16$^{+}$ VLV group exhibited a marked loss of proliferative peaks and overall reduction in B16-F10 cell counts, whereas EO771 proliferation was minimally affected. **(B)** Splenocyte viability. Live-cell gating of CFSE$^{+}$ splenocytes (Pacific Blue-H) at 48 h revealed improved viability in responder cells from B16$^{+}$ VLV-vaccinated mice, consistent with enhanced effector/memory survival. Data are representative of two to four biological replicates, each performed in triplicate wells.

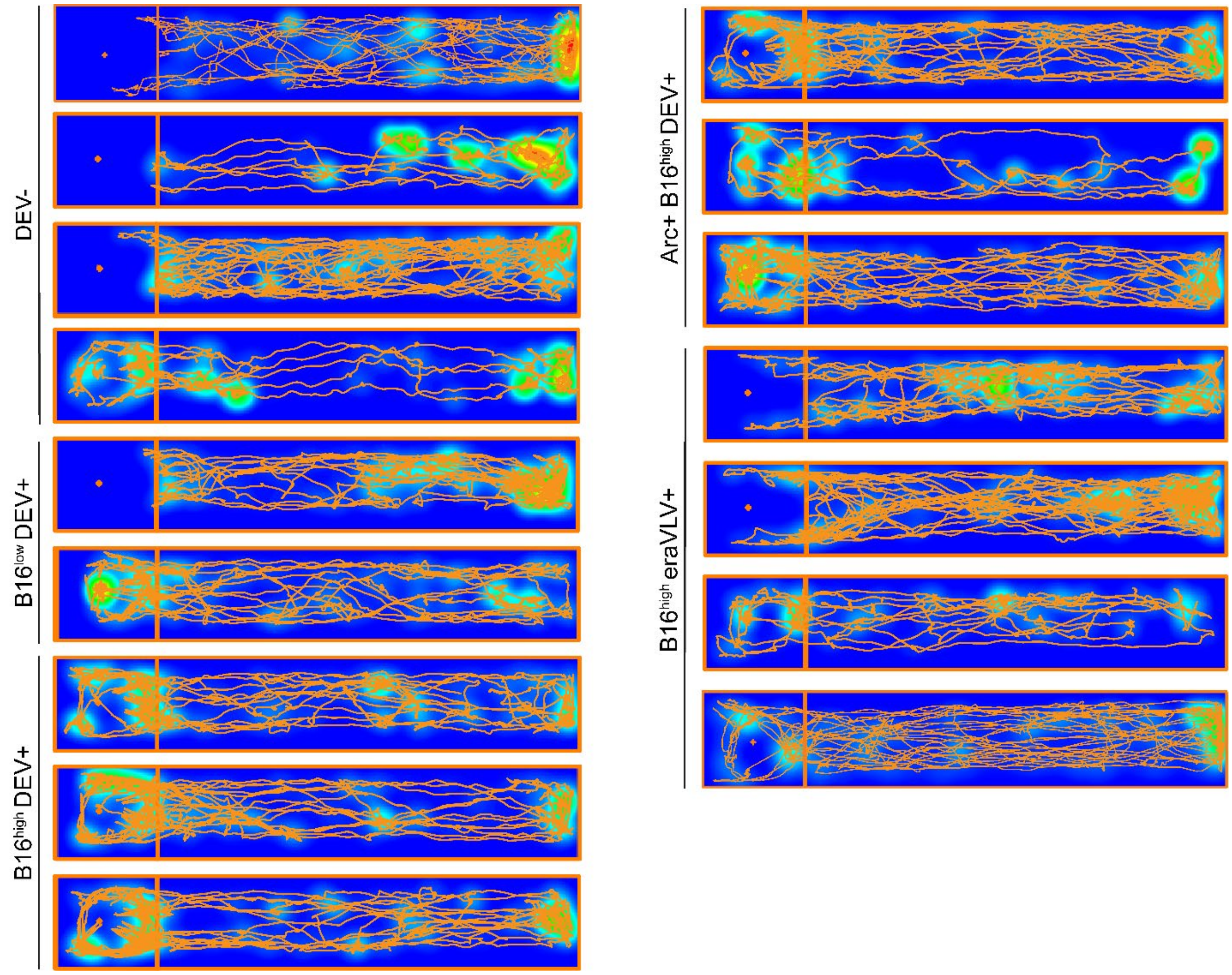


**Figure S13.** *Locomotion and cognitive assessment using a novel object test.* This supplementary figure shows detailed movement tracks of mice from different experimental groups within the novel object test chamber across independent experiments. Orange tracks indicate the paths taken by each mouse, and the accompanying heat maps depict movement density, highlighting regions where mice spent the most time. Besides the negative control mice (newly tumor-challenged), all subjects were vaccinated survivors that withstood two rounds of tumor challenge. Overall, treated survivors performed similarly in this test.

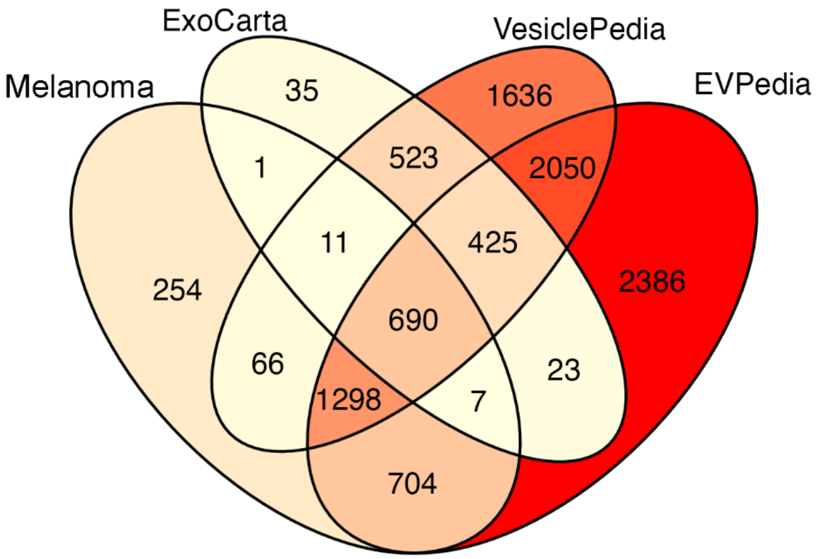


**Figure S14.** *Venn diagram compares the proteins identified in our samples with those annotated in the ExoCarta, Vesiclepedia, and EVpedia databases.*

**Extended methods**

**Cells and cell culture**

Stable cell line culture: DC2.4 murine dendritic cells (Millipore, Cat# SCC142) and B16-F10 melanoma cells (ATCC, CRL-6475) were maintained in DMEM or RPMI-1640 supplemented with 10% FBS (Gibco) at 37 °C in a 5% $CO_2$ incubator. EO771 breast cancer cells (gift from the Weiss Lab, Cornell University) were cultured under identical conditions. Cells were maintained according to ATCC guidelines. No antibiotics were used in stable cell line cultures. For *in vivo* tests, we matched donor strains of cells with corresponding recipient mice (DC2.4 with C57BL/6J for example).

Primary BM GM-CSF/IL-4 cultures (BM-DC/MΦs): Primary bone marrow cells were extracted from wildtype mice (C57BL/6J) after disinfection with 70% ethanol. The femurs and tibiae were dissected and soaked in HBSS (Hank's Balanced Salt Solution, Gibco) or RPMI-1640 medium (ATCC) supplemented with 1% FBS (Gibco). Both ends of the bone were cut open with surgical scissors, and a 25G needle (with a 20-mL syringe) was inserted into the bone cavity to rinse the BM cells out of the femur, whereas a 27G needle was used for the tibia. A total volume of 20 mL complete RPMI-1640 medium was used to slowly (dropwise) flush out BM cells from each femur and a total volume of 10mL was used for each tibia. The cell suspension was passed through 70 µm cell strainer and centrifuged at 180X g for 10 min. The cell pellet was resuspended with ice cold eBioscience red blood cell (RBC) lysis buffer and incubated on ice for 5 minutes to lyse RBCs. Following a second centrifugation, the supernatant was discarded, and the pelleted cells were rinsed with complete medium and collected. $10 \times 10^6$ collected cells were cultured in each tissue culture-treated T25 flask in 5 ml of complete medium (ATCC RPMI-1640 supplemented with 2-mercaptoethanol (Invitrogen), 10% fetal bovine serum (Gibco), Pen-Strep, murine GM-CSF (20 ng/ml, Peprotech) and IL-4 (5 ng/ml, Peprotech). Half of the medium was removed on day 2 and replaced by fresh warm medium supplemented with 40 ng/ml GMCSF and 10 ng/ml IL-4. The culture medium was entirely discarded on day 3 and replaced by fresh warm medium with 20 ng/ml GM-CSF and 5 ng/ml IL-4. On day

6, all cells in the culture including adherent, suspension, and loosely attached cells, harvested by gentle washing with warm DPBS, were pooled and used as the source of leukocyte EVs.

**EV production**

Donor cells underwent transfection, followed by EV isolation and characterization. Thorough titration and time-lapse experiments optimized transfection and EV production. Transfection methods: lipofectamine mMessenger Max RNA transfection. was used to transfect immortal and primary cells. Monitoring transfection efficacy: In addition to stabilizing Arc VLVs, A5U-GFP also served as a fluorescent reporter to closely track transfection efficacy within donor cells. Optimal production time and consistency: Depending on donor cell confluency, the time to achieve peak transfection efficacy and thus EV production can vary. Given the inherent variability in primary cells across different batches, we utilized live cell imaging to monitor GFP expression in donor cells. EVs were produced in a serum-free OptiMEM culture medium to enhance the yield of production. EVs were freshly prepared before both *in vivo* and *in vitro* experiments, as long-term storage was found to reduce their integrity and functionality.

**Fluorescent labeling of EVs**

Freshly isolated EVs ($1 \times 10^{11}$ particles $mL^{-1}$, determined by NTA) were stained with CellMask™ Deep Red (Thermo Fisher Scientific) diluted 1:5000 in sterile, 0.1 µm-filtered DPBS. Reactions were mixed gently and incubated 10 min at room temperature on a rocker, protected from light. Unbound dye was removed by five to six ultrafiltration washes (100 kDa cutoff, 4 °C) with filtered DPBS until the mock DPBS control, processed through the same staining and washing protocol, appeared completely clear under the same fluorescence detection settings. If, after extensive washing, the DPBS blank still exhibits detectable fluorescence above threshold, this indicates that dye micelles or aggregates have formed in the preparation. Such samples should be discarded and re-stained with fresh dye at lower concentration or higher EV density. Washed EVs were resuspended at $1 \times 10^{12}$ particles $mL^{-1}$ for injection. All subsequent handling was performed in low-bind tubes at 4 °C to minimize adsorption and photobleaching.

**Fluorescent NTA:**

The NanoSight NS300 detects nanovesicles with stable fluorophores using a 488 nm or 532 nm laser. We apply green and red filters to spot fluorescent signals. Negative control helps adjust the camera to exclude non-fluorescent signals. In fluorescent mode, individual particle movement helps determine particle size and concentration, as larger particles move slower than smaller ones. In light scatter mode, we measure the total particle count. We use specific fluorescent labels to identify different types of EVs, including general EV markers, Arc, or plasma membrane dyes.

**Mouse experiments and vaccination procedure**

All animal procedures were performed at the Cornell University Center for Animal Resources and Education (CARE) in accordance with institutional ethical guidelines and approved IACUC protocols #2020-0037 and #2023-0101. Mice were housed under specific pathogen-free conditions (12-h light/dark cycle, 22 ± 1 °C, 40–60% humidity) with ad libitum access to food and water. Wild-type C57BL/6J mice (Jackson Laboratory) aged 6-10 weeks of both sexes were randomly assigned to treatment groups. Each cohort contained littermates balanced by sex to minimize batch variability.

<u>Vaccine administration:</u> Engineered vesicle vaccines ($B16^{+}$ VLVs or DEVs) were injected intravenously via the retro-orbital sinus once per week for four consecutive weeks. To minimize local irritation, injections alternated between the left and right eye each week. All vesicle preparations were produced fresh before dosing and quantified by NTA. Each dose consisted of 50 µL of EV suspension containing $6 \times 10^{10}$ particles per mouse, corresponding to the empirically determined minimal effective dose. The exact particle concentrations and dosing records for each batch are provided in Supplementary Table 1. Control mice received equivalent volumes of sterile DPBS.

<u>Post-injection monitoring:</u> Mice were observed until full recovery from anesthesia and monitored daily for signs of distress, weight loss, or injection-site inflammation. Body weight, activity, and grooming were

recorded weekly. No adverse effects or mortality were observed in any treatment group under the dosing regimen used.

**Microscopy**

Epifluorescence live-cell imaging: Cells were plated on tissue-culture–treated glass-bottom dishes and imaged on a Cytation 7 multimode imager (BioTek) equipped with temperature and $CO_2$ control. Images were acquired at multiple stages of transfection and culture to monitor cell morphology and transfection efficiency. DAPI counterstaining was used to quantify nuclei and assess cell density and brightness relative to unstained controls. Acquisition and analysis were performed using Gen5 software. DNA and RNA transfection parameters were optimized in real time based on GFP/A5U-GFP fluorescence intensity and cell viability. Vesicle production was subsequently quantified by nanoparticle tracking analysis (NTA), and EV suspensions were diluted according to the desired particle dose.

Confocal microscopy: High-resolution imaging was performed using a Zeiss LSM 980 confocal laser-scanning microscope equipped with 405-, 488-, 561-, and 640-nm laser lines and multiple objectives (5× and 20× air, 10× water immersion, 60× oil immersion). Optical sections were acquired with pinhole set to 1 Airy unit and frame averaging (2-4×) to improve signal-to-noise. Images were processed using ZEN blue and analyzed in ImageJ (NIH) for fluorescence intensity profiles, co-localization, and vesicle distribution.

Immunocytochemistry (ICC) and Immunohistochemistry (IHC): For ICC, adherent cells were fixed with 4% paraformaldehyde (PFA) for 15 min, permeabilized with 0.1% Triton X-100, and blocked in 3% normal goat serum before incubation with primary antibodies (typically overnight at 4 °C). For IHC, cryosections (10-30 µm) or paraffin-embedded tissue sections were rehydrated and subjected to antigen retrieval when necessary. Primary antibodies against EV or cellular markers were detected with fluorophore-conjugated secondary antibodies. Samples were mounted in ProLong™ Gold Antifade Mountant with DAPI (for thick tissue) or PVA DABCO (for thin sections). Imaging parameters were matched across samples to allow quantitative comparison of staining intensity.

In vivo fluorescence imaging (IVIS Spectrum): Mice were perfused trans-cardially with PBS under deep anesthesia to reduce background fluorescence before organ harvest. Whole-organ fluorescence was imaged on a Perkin Elmer IVIS Spectrum system using appropriate excitation/emission filters for CMDR. Negative (unlabeled) controls were included in every session to correct for tissue autofluorescence. Quantification of radiance (photons $s^{-1}$ $cm^{-2}$ $sr^{-1}$) was performed with Living Image or Aura software, and region-of-interest values were exported to ImageJ for further normalization and statistical analysis.

**Tumor measurement and imaging** Manual tumor size measurement: Tumor measurements commenced five days after the injection of B16-F10 melanoma cells. Measurements were continued daily thereafter to capture the progressive changes in tumor size throughout the experimental period, using digital calipers. To minimize bias and ensure the reliability of data, tumor measurements were performed in a double-blind manner by a group of five researchers (independent measurements). This approach involved concealing the treatment groups from the individuals performing the measurements, thereby preventing potential observer bias. Micro-CT scan: Micro-computed tomography (micro-CT) scans were conducted to enable non-invasive and high-resolution imaging of tumor growth in live animals. Prior to scanning, animals were anesthetized to minimize motion artifacts during image acquisition. Micro-CT scans were performed using specialized imaging equipment, which generated cross-sectional images of the entire body, including tumor-bearing regions. The acquired micro-CT images were subjected to detailed analysis using dedicated imaging software. This software allowed for the precise delineation and measurement of tumor volumes based on differences in tissue density. Tumor volumes were quantified by outlining the tumor boundaries on each cross-sectional image and summing the volumes across all slices. This rigorous analysis facilitated accurate assessment of tumor growth kinetics and treatment efficacy over time.

**Analysis of immune response**

Cell collection: Following treatment, mice were euthanized, and spleens and lymph nodes were collected aseptically. Splenocytes were isolated by mechanically disrupting the spleen tissue and passing it through a cell strainer (70 μm). Red blood cells were lysed using the RBC lysis buffer, and the remaining splenocytes were washed and resuspended in optiMEM or flow cytometry buffer for downstream analysis. Lymphocytes were harvested from lymph nodes by mechanical disruption and filtration through cell strainers (70 µm and then 40 µm) to obtain a single-cell suspension. The cells were then washed and resuspended in a suitable buffer for subsequent analysis. Flow cytometry: Prior to analysis, cells were stained with a panel of fluorescently labeled antibodies targeting specific cell surface markers. In addition to CD45 and live/dead stain, our panel included antibodies against CD3, CD4, and CD8, which are commonly used to identify T cell subsets. Additionally, other markers relevant to T cell proliferation and activation were included in the staining panel to provide comprehensive characterization of the immune response, including CD44, CD62L, INFgamma. Stained cells were analyzed using a flow cytometer (Novocyte quanteon 4025), which detects and quantifies the fluorescence emitted by individual cells as they pass through a laser beam. Data acquisition was performed using specialized Novocyte software, with the data analyzed using FlowJo to categorize the cell populations.

ELISA: At time points of 3, 4, 6, 9, and 13 weeks after the first dose, blood samples were collected by completing cheek blood drawings. These samples were analyzed using enzyme-linked immunosorbent assays (ELISAs) to determine the protein levels of relevant antibodies (IgG, IgM, and IgA). This involves coating 96 well plates with Ova protein, blocking them with a blocking solution (10% NGS in SuperBlock), adding samples at various concentrations, adding secondary antibody, developing the plates using TMB substrate, stopping the reaction using sulfuric acid, and imaging the plates on a Cytation imaging machine.

qPCR: Total RNA was extracted from harvested cells using Trizol/BCP followed by DNase treatment and RNA purification (PureLink, Invitrogen). cDNA synthesis: 2 µg of total RNA was then reverse transcribed into cDNA using *Applied Biosystem* High-Capacity cDNA *Reverse Transcription* Kit. Primers: Resulting cDNA was diluted and used as the template for qPCR using PowerUp SYBRgreen Master Mix (Thermo Fisher Scientific) with primers against various cytokines and virus-like components. The list of qPCR primers can be found in supplementary table 2. The specificity of primers was confirmed by a melt curve analysis. The qPCR thermal cycling conditions for PowerUp™ SYBR™ Green Master Mix start with an initial denaturation at 95°C for 2 minutes, followed by 40 cycles of 95°C for 15 seconds (denaturation) and 60°C for 1 minute (annealing/extension). For data analysis, either the ΔCt method (normalizing target gene expression to a reference gene within a single sample) or the ΔΔCt method (comparing normalized expression between samples, such as treated *vs.* untreated) was applied.

ELISpot assay: Following the booster dose administration, an Enzyme-Linked Immunospot (ELISpot) assay was performed to assess T cell activation and memory response elicited by the treatment. 96-well plates were coated with antibodies specific to IFN-γ. Treated cells were then added to the wells and incubated under conditions conducive to cytokine secretion (Ova MHC-I or MHC-II peptides, or B16-F10 EV coincubation). Upon cytokine secretion by activated T cells, cytokine molecules were captured by the immobilized antibodies, resulting in the formation of visible spots at the locations of cytokine-secreting cells. After incubation, the plates were washed to remove unbound cells and cytokines, followed by the addition of detection antibodies and a substrate solution. This led to the development of colored spots corresponding to individual cytokine-secreting cells. The number of spots in each well, representing the frequency of cytokine-secreting cells, was quantified using Cytation 7 plate imager. The magnitude and quality of T cell responses were assessed based on the frequency and distribution of cytokine-secreting cells.

***Ex vivo* cytotoxicity and splenocyte-tumor co-culture assay**

Generation of donor dendritic cells and vesicle vaccines: BMDCs were isolated from WT and $Arc^{-/-}$ mice as described above. One subset of WT BMDCs was transfected with Arc and A5U mRNAs using Lipofectamine MessengerMAX in Opti-MEM for 6 h to generate Arc-overexpressing donor cells, while the other WT and $Arc^{-/-}$ BMDCs were left untransfected. Each donor group was subsequently exposed for 24 h to B16-F10 melanoma-derived EVs serving as tumor antigen sources for DEV vaccine production. After removal of residual tumor EVs and cell debris, DEVs were purified and quantified by NTA.

**Vaccination and splenocyte collection**

C57BL/6J mice were vaccinated intravenously with vesicles generated from the three donor groups: (1) $Arc^{+}$ $A5U^{+}$ BMDCs, (2) untransfected WT BMDCs (endogenous $mArc^{+}$), and (3) $Arc^{-}/^{-}$ BMDCs (mArc-null control). Eight days post-vaccination, mice were euthanized under isoflurane anesthesia, and spleens were aseptically harvested. Single-cell suspensions were prepared by mechanical disruption through a 70 μm cell strainer, followed by red blood cell lysis (ACK buffer, 5 min, room temperature). Splenocytes were washed twice in Opti-MEM and resuspended at $6 \times 10^{6}$ cells $mL^{-1}$.

<u>Labeling of effector and target cells:</u> Splenocytes (effectors) were labeled with CFSE (Thermo Fisher, 5 μM in PBS, 10 min, 37 °C), quenched with FBS, and washed twice. Target B16-F10 melanoma cells were pre-stained with NucSpot™ 650 nuclear dye (Biotium; 1:1000 dilution, 10 min, RT) to enable discrimination in co-culture.

<u>Co-culture conditions:</u> Effector and target cells were co-incubated in 96-well flat-bottom plates at an E:T of ~200:1 ($0.6 \times 10^{6}$ splenocytes + $3 \times 10^{3}$ tumor cells per well) in 100 μL serum-free Opti-MEM. To maintain splenocyte viability without driving nonspecific activation, cultures received PHA-P at 0.5 μg $mL^{-1}$ (low dose). Plates were incubated 24-48 h at 37 °C, 5% $CO_2$, without agitation to preserve effector-target conjugates. Controls included targets only, effectors only, and splenocytes from PBS-injected mice (±PHA-P), plus medium-only wells to monitor background.

<u>Imaging and analysis:</u> After 24 h, the culture supernatant containing nonadherent splenocytes was gently removed, and wells were washed twice with pre-warmed Opti-MEM to remove unbound cells. Adherent tumor cells were immediately imaged by epifluorescence microscopy (Cytation 7, BioTek) using GFP (for CFSE) and Cy5 (for NucSpot) channels to visualize effector-target interactions. Interaction frequency was quantified as the percentage of tumor cells in direct contact with $CFSE^{+}$ splenocytes per high-power field (five random fields per well).

All assays were performed in triplicate from independent mice. Results were validated by comparing $Arc^{+}$ DEV-, WT DEV-, and $Arc^{-}/^{-}$ DEV–vaccinated splenocytes for relative cytotoxic capacity and effector–target conjugate formation.

**Mass spectrometry analysis**

Protein Isolation: Proteins associated with Arc VLVs and control DEVs were isolated from the culture supernatant of genetically modified DCs overexpressing the Arc protein and control DCs. The culture supernatant was collected and subjected to centrifugation to remove cellular debris, followed by TFF and ultrafiltration to isolate and concentrate the EVs. The isolated EVs were lysed using an IP lysis buffer containing detergents and protease inhibitors. This lysis step involved one freeze-thaw cycle and vortexes while incubating on ice to disrupt the EV membrane, releasing the protein cargo into solution. The lysate was then centrifuged to remove insoluble debris, and the supernatant containing solubilized proteins was collected for downstream processing. The concentration of solubilized proteins was determined using a protein quantification assay (Pierce 660). This step ensured that equal amounts of protein were loaded onto the subsequent steps of the analysis, enabling accurate and reproducible results. To prepare the proteins for electrophoresis, the solubilized protein sample was denatured and reduced. Denaturation involved heating the sample in the presence of denaturing agents sodium dodecyl sulfate (SDS), which disrupted the protein's secondary and tertiary structures, while reduction involved the addition of reducing agents (5% beta-mercaptoethanol) to break disulfide bonds.

SDS-PAGE Electrophoresis: The denatured and reduced proteins were separated by size using sodium dodecyl sulfate-polyacrylamide gel electrophoresis (SDS-PAGE). The protein sample was loaded into wells of an acrylamide gel and subjected to an electric field, causing the proteins to migrate through the gel matrix based on their molecular weight. This step allowed for the separation of proteins according to their size. Following electrophoresis, the protein bands were visualized using staining methods (SimplyBlue or silver staining). These staining techniques allowed for the detection of protein bands within the gel, which appeared as distinct bands against a clear background.

In-gel trypsin digestion: The gel bands from the lanes of interest were excised, cut into ~2 mm cubes and subjected to in-gel digestion. The excised gel pieces were washed/incubated at room temperature consecutively with 150-400μL deionized water for 5 minutes, followed by 150-400 μl 50mM ammonium bicarbonate in water/50% acetonitrile (ACN) for 10 minutes and finally 150-400µl 100% ACN for 5 minutes. The dehydrated gel pieces were dried in a speed vacuum (SpeedVac SC110 Thermo Savant, Milford, MA) and reduced with 50-250µL of 10mM DTT (dithiothreitol) (w/v) **i**n 100mM ammonium bicarbonate in water for 1 hour at 60°C, then alkylated by adding 50-250µL of 55mM iodoacetamide in 100mM ammonium bicarbonate (w/v) and incubation at room temperature, in the dark, for 45 minutes. Wash steps were repeated as described above. The gel pieces were dried in a speed vacuum and rehydrated with 40 - 120 µl trypsin (Promega Sequencing Grade) at 10ng/µl (w/v) in 50mM ammonium bicarbonate/10% ACN on ice for 20 minutes, topped with 10-50 µl 50 mM ammonium bicarbonate in water, and incubated at 37°C for 16 hours. The digestion was stopped by addition of 50-200 µl 2% formic acid (FA) in water, incubated at room temperature for 10 minutes and the supernatant transferred to a clean polypropylene low-bind microfuge tube. The gel pieces were further extracted twice by adding 100-400µl of 50% ACN/5% FA and vortexing at 1500 rpm for 10 minutes followed by sonication for 5 minutes, and once by adding 100-400µl of 90% ACN/5% FA with incubation at room temperature for 5 minutes. All supernatants were combined in the corresponding microfuge tube, dried in a speed vacuum, and redried from 100 µl water. The final sample was reconstituted in 2% ACN/0.5% FA and filtered through a 0.22µm cellulose acetate spin filter (Corning Costar Spin-X) prior to nanoLC-MS/MS analysis.

<u>Protein identification by nano LC/MS/MS analysis:</u> The analysis was carried out using an Orbitrap Fusion$^{TM}$ Tribrid$^{TM}$ (Thermo-Fisher Scientific, San Jose, CA) mass spectrometer equipped with a nanospray Flex Ion Source, and coupled with a Dionex UltiMate 3000 RSLCnano system (Thermo, Sunnyvale, CA)(Yang et al., 2018). The peptide samples (10 µL) were injected onto a PepMap C-18 RP viper trapping column (5 µm, 100 µm i.d x 20 mm) at 20 µL/min flow rate for rapid sample loading and then separated on a PepMap C-18 RP nano column (2 µm, 75 µm x 25 cm) at 35 °C. The tryptic peptides were eluted in a 90-min gradient of 5% to 35% ACN in 0.1% formic acid at 300 nL/min, followed by an 8-min ramping to 90% ACN-0.1% FA and an 8-min hold at 90% ACN-0.1% FA. The column was re-equilibrated with 0.1% FA for 25 min prior to the next run. The Orbitrap Fusion was operated in positive ion mode with spray voltage

set at 1.2 kV and source temperature at 275°C. External calibration for FT, IT and quadrupole mass analyzers was performed. In data-dependent acquisition (DDA) analysis, the instrument was operated using FT mass analyzer in MS scan to select precursor ions followed by 3 second “Top Speed” data-dependent CID ion trap MS/MS scans at 1.6 m/z quadrupole isolation for precursor peptides with multiple charged ions above a threshold ion count of 10,000 and normalized collision energy of 30%. MS survey scans at a resolving power of 120,000 (fwhm at m/z 200), for the mass range of m/z 300-1600. Dynamic exclusion parameters were set at 50 s of exclusion duration with ±10 ppm exclusion mass width. All data were acquired under Xcalibur 4.4 operation software (Thermo-Fisher Scientific).

Data analysis: The DDA raw files with MS and MS/MS were subjected to database searches using Proteome Discoverer (PD) 2.4 software (Thermo Fisher Scientific, Bremen, Germany) with the Sequest HT algorithm. The PD 2.4 processing workflow containing an additional node of Minora Feature Detector for precursor ion-based quantification was used for protein identification and relative quantitation of identified peptides and their modified forms. The database search was conducted against *Mus musculus* NCBI ref database which contains 28230 sequences and the EVs database with added chicken Ova customer shared with us which contains 74 sequences. The peptide precursor tolerance was set to 10 ppm and fragment ion tolerance was set to 0.6 Da. Oxidation of M, deamidation of N and Q were specified as dynamic modifications of amino acid residues; protein N-terminal acetylation, M-loss and M-loss plus acetylation were set as a variable modification; carbamidomethyl C was specified as a static modification. Only high confidence peptides defined by Sequest HT with a 1% FDR by Percolator were considered for confident peptide identification. Relative quantitation of identified proteins between the paired groups was determined by the Label Free Quantitation (LFQ) workflow in PD 2.4. After retention time alignment for each of the identified peptides across samples, the precursor abundance intensity for each peptide identified by MS/MS in each sample were automatically determined and their unique and razor peptides for each protein in each sample were summed and used for calculating the protein abundance by PD 2.5 software.

LC-MS/MS Analysis: A NanoElute LC system coupled to a timsTOF Pro (Bruker Daltonics, Germany) via a CaptiveSpray source was employed for the analysis. Samples (100 ng) were injected onto an in-house packed column (75 mm x 15 cm, 1.9 μm ReproSil-Pur C18 particles, Dr. Maisch GmbH, Germany) maintained at 40 °C. The mobile phases consisted of buffer A (0.1% formic acid in water) and buffer B (0.1% formic acid in acetonitrile). A 21-minute gradient was applied, starting with 2% buffer B and increasing to 30% over 17.8 minutes, followed by a rapid increase to 95% buffer B by 18.3 minutes, and held for an additional 2.4 minutes. Mass spectrometry data were acquired using a data-independent acquisition parallel accumulation-serial fragmentation (diaPASEF) method with 16 m/z and ion mobility windows. The electrospray voltage was set at 1.5 kV, with the ion transfer tube maintained at 180 °C. Full MS scans were conducted over an m/z range of 100-1700. Collision energy was linearly ramped from 20 eV at $1/K0 = 0.6\ V{\cdot}s/cm^2$ to 59 eV at $1/K0 = 1.6\ V{\cdot}s/cm^2$. Data were processed using DIANN version 1.8 software (Demichev et al., 2020), with default settings for peptide and protein identification and quantification from diaPASEF data. An in-house spectral library was utilized, generated from the UniProt-SwissProt Homo sapiens database (Taxon ID 9606, downloaded on 01/20/2023, containing 20,404 entries). Cysteine carbamidomethylation was specified as a fixed modification, while methionine oxidation and acetylation were considered variable modifications. The false discovery rate (FDR) was controlled to be below 1% at both the peptide and protein levels.
